# Rapid and Reliable Structural Modeling of Adaptive Immune Receptors Using an Optimized AlphaFold3 workflow

**DOI:** 10.64898/2025.12.19.695451

**Authors:** Alexandre Jann, Marta A. S. Perez, Vincent Zoete

## Abstract

AlphaFold (AF), a deep-learning based protein modelling approach, has revolutionized structural biology by generally achieving near-experimental accuracy in protein structure prediction. An impactful application of AF is the modelling of antibodies (Abs) and T-cell receptors (TCRs), key mediators of cellular immunity, whose structural specificity underlies responses in cancer, infection, and autoimmune diseases. In this work, we analyse AF3 performance by systematically examining how MSA composition, number of inference phases and inference parameters affect prediction accuracy and computational efficiency. We present an acceleration of ∼45-fold of the AF3 MSA phase using reduced UniRef90 subsets, combined with up to a 3.6-fold increase in AF3 inference speed through optimized parameters. We provide a highly accurate variant of the AF3 workflow specifically optimized for the modelling of the Abs and TCRs receptor domains, enabling rapid, reliable structural predictions at a scale suitable for high-throughput immunological studies. Our findings provide a foundation for faster therapeutic discovery and deeper molecular mechanism understanding of immune recognition.

**Teaser:** By improving key steps in the process, we made AlphaFold3 about 40 times faster at modeling specific immune proteins

## Introduction

The ability to accurately predict protein structures has been revolutionized by AlphaFold (AF)[1], a deep learning (DL) framework that generally demonstrates near-experimental accuracy across most of the human proteome. AlphaFold3 (AF3)[2], released in 2024, is the most recent and advanced version of AF for biomolecular structure prediction and interaction analysis. Unlike AlphaFold2 (AF2)[3], AF3 significantly reduces the reliance on Multiple Sequence Alignment (MSA) representations. While MSA information is still used, its role is greatly diminished in favor of single and pair representations, which helps to simplify the model’s architecture. The single representation refers to per-residue or per-atom embeddings that capture local biochemical properties. The pair representation encodes features between every pair of residues or atoms, capturing spatial proximity, geometric constraints, and co-evolutionary signals. AF3 adopts new components, such as a Pairformer and a Diffusion-based Generative Model, to replace the Evoformer stack used in AF2. These new modules are responsible for updating both pairwise and single representations in a more flexible and generalizable manner, allowing AF3 to support heterogeneous biomolecular assemblies.

Despite the reduced influence of the MSA on model construction in AF3, the MSA component remains the slowest step in the workflow, representing more than 90% of the end-to-end execution time. Consequently, MSA generation imposes a disproportionate runtime and emerges as a bottleneck in AF3, notably for studies requiring the generation of large numbers of 3D structures. Some alternative approaches mitigate the expensive MSA computation, ColabFold[4] uses a different software to compute the MSA faster while ImmuneBuilder[5] removes the need of MSA as an input by training their machine learning model exclusively on immune proteins.

T-cell Receptors (TCRs) and Antibodies (Abs) are key mediators of cellular immunity, whose structural specificity underlies responses in cancer, infection, and autoimmune diseases. TCRs and Abs exhibit exceptional sequence diversity and flexible regions, posing challenges in the creation of 3D models. Abs and TCRs are distinct from other proteins due to their conserved structure in the framework region but large structure variability in the Complementarity-Determining Regions (CDR) regions. Our group develops 3D-based predictors of TCR and Ab repertoires analysis and specificity prediction[6, 7], which rely on the rapid modeling of large numbers of receptor domains.

In this paper, we conducted a comprehensive performance analysis of AF3 for modelling TCRs and Abs. We focus on two phases of the AF3 workflow: (i) the MSA phase, which prepares evolutionary features from homologous sequences, and (ii) the inference phase, which performs structure prediction using deep neural network modules, primarily the Pairformer and Diffusion module, to iteratively update pair representations that capture atomic interaction relationships. By benchmarking end-to-end execution time across multiple AF3 workflow variants, with different MSA databases and number of seeds, we demonstrate that the AF3 efficiency can be improved by using subsets of the UniRef90 database [8] in the MSA phase (**Figure 1a-b)**. On our desktop system, AF3 MSA pipeline execution time is accelerated by up to 45-fold while keeping the TCRs and Abs models at near-experimental accuracy. We also improved AF3 inference parameters to better fit Ab and TCR modelling, reducing the new bottleneck step of model generation. By allowing 9 inferences to run in parallel on a single 24Go GPU and optimizing AF3 specific parameters, we reduced inference time by an additional 1.5 to 3.6-fold depending on input size (**Figure 1a**). Developing specialized AF3 variants that retain AF’s predictive accuracy while making it extremely fast is critical for advancing structural immunology and enabling large-scale modeling of immune repertoires. Our approach could be readily generalized to other protein families with large numbers of experimentally determined 3D structures in the Protein Data Bank (PDB), such as kinases or proteases or MHC complexes.

**Figure 1.**
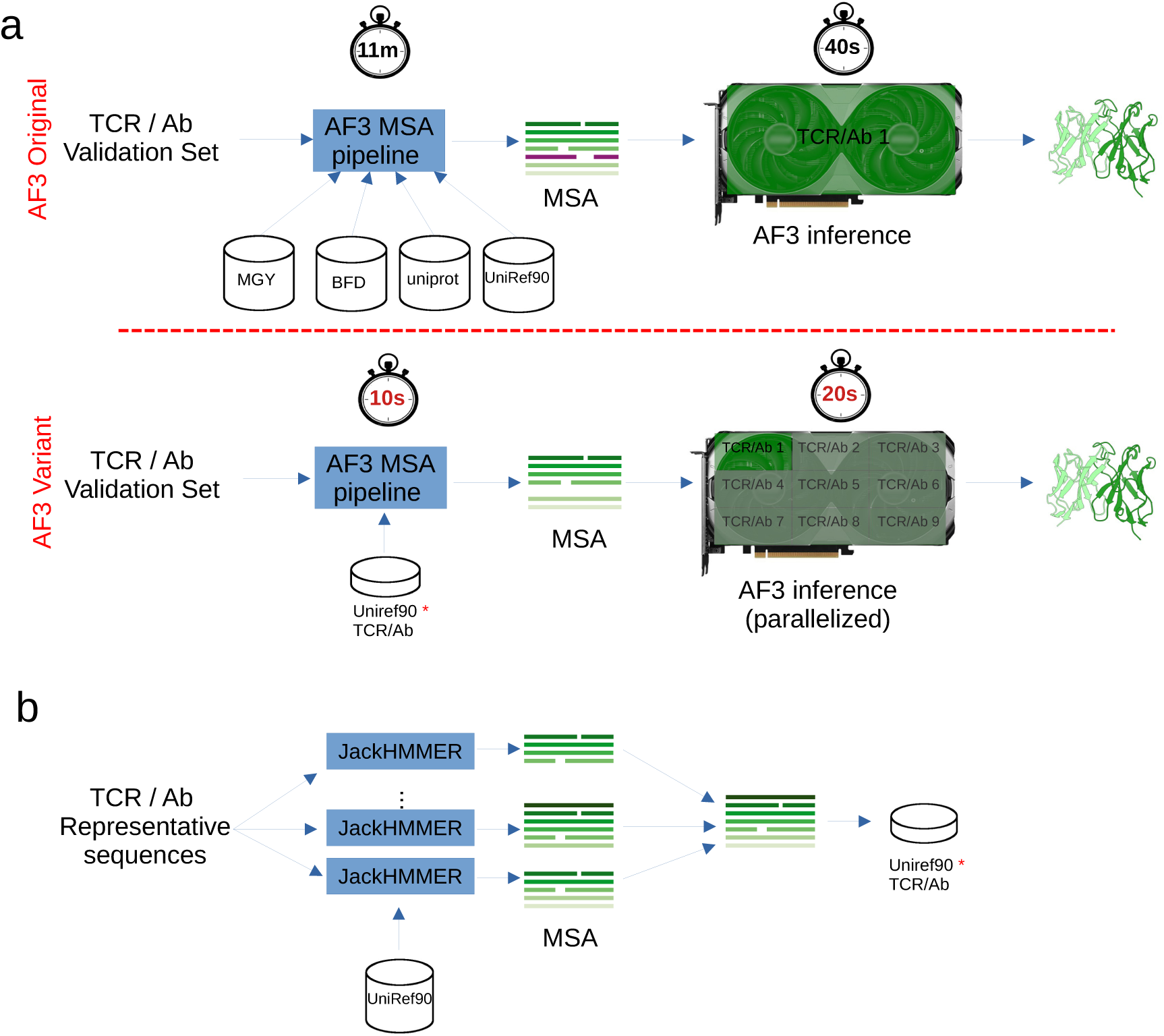
**a**) General overview of AF3 original workflow and our AF3 variant workflow, along with running time of the corresponding AF3 MSA and inference phases. The original AF3 uses 4 databases to build the MSA input while AF3 variant uses a much smaller database named UniRef90-TCR or UniRef90-Ab. **b**) Process used to build the reduced Uniref90-TCR/Ab databases, by sequentially generating MSAs for individual TCRs using JackHMMER and the UniRef90 database, and aggregating the resulting sequences.

## Results

In this results section, we present AF3 workflow variants specifically developed for the modelling of the receptor domain of the Abs and TCRs. We show that the AF3 end-to-end runtime can be substantially reduced by generating a curated subset of UniRef90 sequences, rather than relying on the full databases used by AF3. Using the AF3 ranking score, we further demonstrate that a single seed is sufficient to obtain highly accurate models, eliminating the need for multiple runs. In addition, we accelerate the AF3 inference stage by up to 3.6-fold through optimized parameter settings. Notably, we show that our AF3 workflow variant achieves near-experimental accuracy, even for challenging CDRs such as antibody CDRH3 and T-cell receptor CDR3b, while relying solely on the reduced database and a single seed.

### Less than 3% of the UniRef90 sequences are used in AF3 MSA for TCR and Ab modelling

AF uses multiple sequence alignments (MSAs) computed by JackHMMER[9] to extract information from evolutionarily related sequences corresponding to the input proteins. Homologous sequences are retrieved for each protein chain in a very time-consuming process, reading sequences from huge databases such as UniRef90, which contains 153,742,194 sequences, to ensure general applicability.

To evaluate the Uniref90 sequences selected by AF3 for the modelling of TCRs we used a non-redundant set of 3’213 TCRs from the VDJ database [10, 11] (**Methods**). By sequentially generating MSAs for individual TCRs and aggregating the resulting sequences, we found that the size of the cumulative sequence database after treating 500 TCRs (15% of the number of samples used) already contains more than 95% of the total number of sequences, becoming sufficient to reproduce the UniRef90-based MSAs of all subsequent TCRs (**Figure 2 a-c**). Across the entire VDJ set, 261,055 sequences are used for TCRα (**Figure 2a** and **2g**), 208,047 for TCRβ (**Figure 2b** and **2g)**, and 261,780 for TCRαβ (**Figure 2 c)** and **2g**), indicating that the α chain incorporates more UniRef90 sequences than the β chain, with substantial overlap between the sequences used for both chains. Less than 3% of the UniRef90 sequences are used in the MSA for the entire set suggesting that the size of the reference database used for MSA could be substantially reduced if AF3 is applied to this specific protein family. These sequences were compiled into a new database called UniRef-TCR. Among the UniRef90 codes present in the reduced database UniRef-TCR and successfully mapped to a protein domain or family using the UniProt ID mapping tool, (https://www.uniprot.org/id-mapping), 98.7% were assigned to proteins with known gene ontology relate to the immune system (i.e. adaptive immune response [GO:0002250]) or with immune-related protein name (i.e. Ig-like domain-containing protein)..

**Figure 2.**
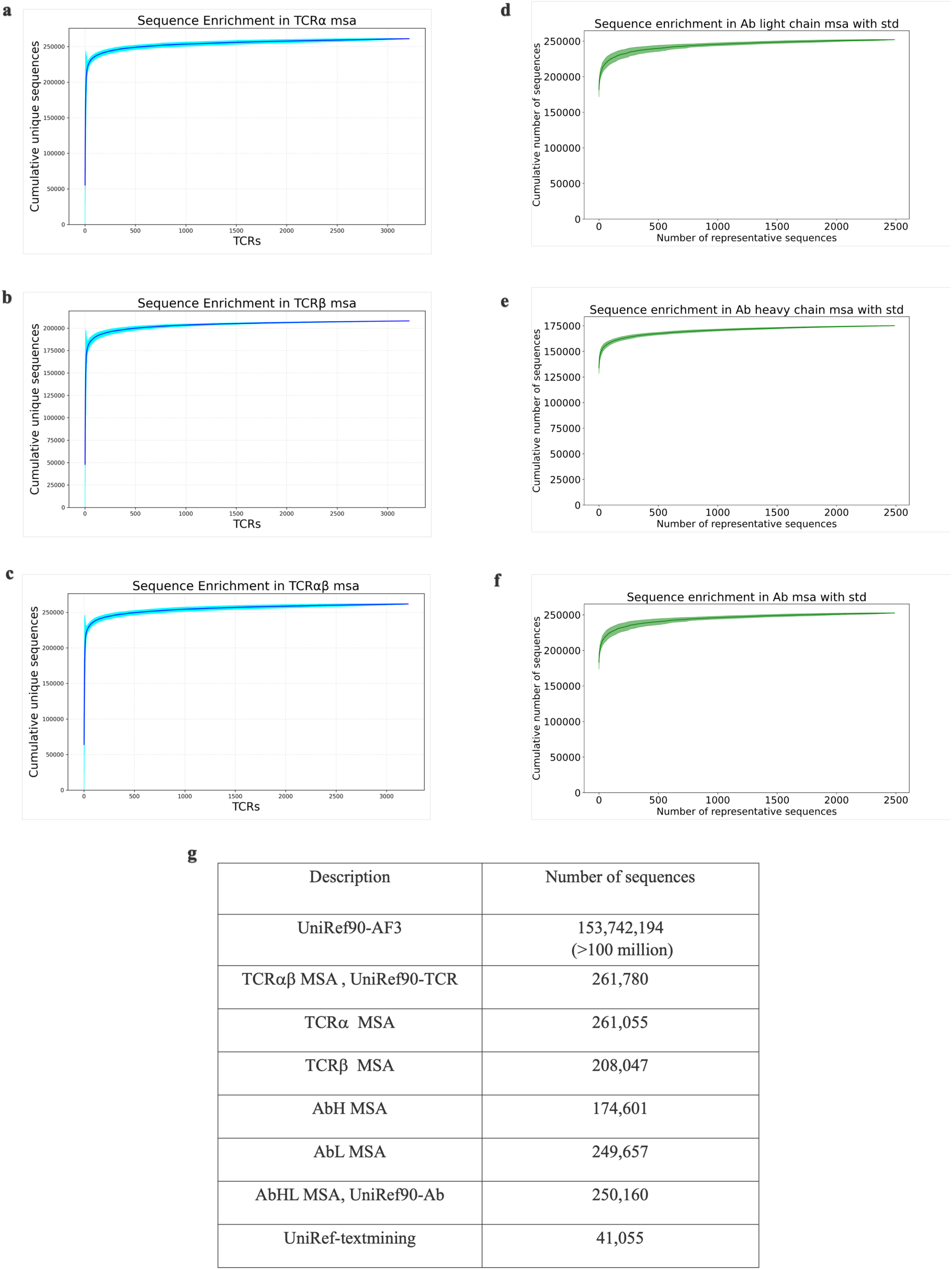
Convergence of the aggregated MSA-derived sequence databases. (**a-c**) Cumulative number of unique sequences obtained from UniRef90 by generating multiple sequence alignments (MSAs) for individual T cell receptors (TCRs) and aggregating the resulting sequences, plotted as a function of the number of TCRs processed, for a set 3,213 unique TCRs from VDJ database. The convergence is shown over 500 random shuffling of the order in which the TCRs were processed. **a)** and **b)** show analyses for TCR α chains and β chains, respectively, and **c**) present analyses of paired αβ TCRs. (**d-f**) Sequence accumulation obtained from the sequential MSA analysis of a set of 2,490 reference Abs, and respective standard deviation computed over 100 random shuffling of the order in which the antibodies were processed. **d)** corresponds to the Ab’s heavy chain MSA analysis, **e)** to Ab’s light chain, **f)** Ab’s heavy and light chain together and **g)**. Number of UniRef90 sequences in the AF3 Uniref90 database, number of sequences from UniRef90 used either in the MSA of TCRa, TCRb and TCRab paired for a set of 3,213 unique TCRs from VDJ database or used in the MSA of AbH, AbL and AbHL paired for a set of 2,490 Abs. Number of sequences in a non-exhaustive database containing just chains belonging to TCRs and Abs.

We also performed the same exercise for a set of 2’490 reference Abs and aggregated a total of 174,601 unique sequences from the UniRef90 database after sequentially treating the heavy chain, 249,657 for the light chain and 250,160 for both heavy and light chains (**Figure 2g**). Similarly to TCR sequences, 98.6% of UniRef90 cluster codes successfully mapped to a Uniprot protein with a gene ontology or protein name related to adaptive immunity. These sequences were aggregated into a new database called UniRef-Ab. The convergence of sequence accumulation during the sequential processing of antibodies, which supports the sufficiency of the final sequence datasets, could in principle depend on the order in which antibodies are analyzed. To assess if convergence is robust to Abs processing order, we performed 100 accumulation runs, each initiated from a different random shuffling of the reference antibody set. The resulting standard deviation of the cumulative number of sequences as a function of the number of reference Abs processed is shown in **Figure 2 d–f**. These results demonstrate that convergence is independent of antibody processing order, supporting the comprehensiveness of the final sequence datasets. We expect this behavior to extend to the construction of the TCR datasets as well. Notably, the accumulation analysis (**Figure 2 d–f**) reveals that, on average, approximately 80% of all unique sequences are already captured after processing the first five reference Abs, indicating substantial sequence sharing across multiple Abs.

Another new database, UniRef-TextMining, containing 41,055 sequences, was generated by text-mining the UniRef90 database for terms associated with TCRs or Abs.

These specialized databases, UniRef-TCR, UniRef-Ab and UniRef-textminining were used as the only databases to generate MSA instead of the databases usually used during a normal AF3 run. The four databases used by AF3 to gather homologous sequences for building the MSA are (i) UniRef90, a clustered version of UniProt with sequences merged at 90 % identity for broad evolutionary coverage (described as UniRef90-AF3 in **Figure 2g**, (ii) MGnigy, large metagenomic sequence database that greatly increases depth of evolutionary information, (iii) BFD (Big Fantastic Database), one of the largest collection of protein families and (iv) the full UniProt protein sequence database[3]. AF3 can also generate MSAs for RNA chains using nucleotide resources (not described because were not used).

### Focused sequence databases can be used for Ab and TCR modelling while keeping accuracy near experimental resolution

To evaluate the impact of reducing the reference database used for MSA, we assessed model performance and accuracy across different AF3 workflow variants that incorporate sequence databases of varying sizes. Model accuracy was quantified by calculating the root mean square deviation (RMSD) between the predicted 3D structure generated with a given AF3 variant (both the original and the reduced-database versions) and the corresponding experimental PDB structure. **Figure 3**.

**Figure 3.**
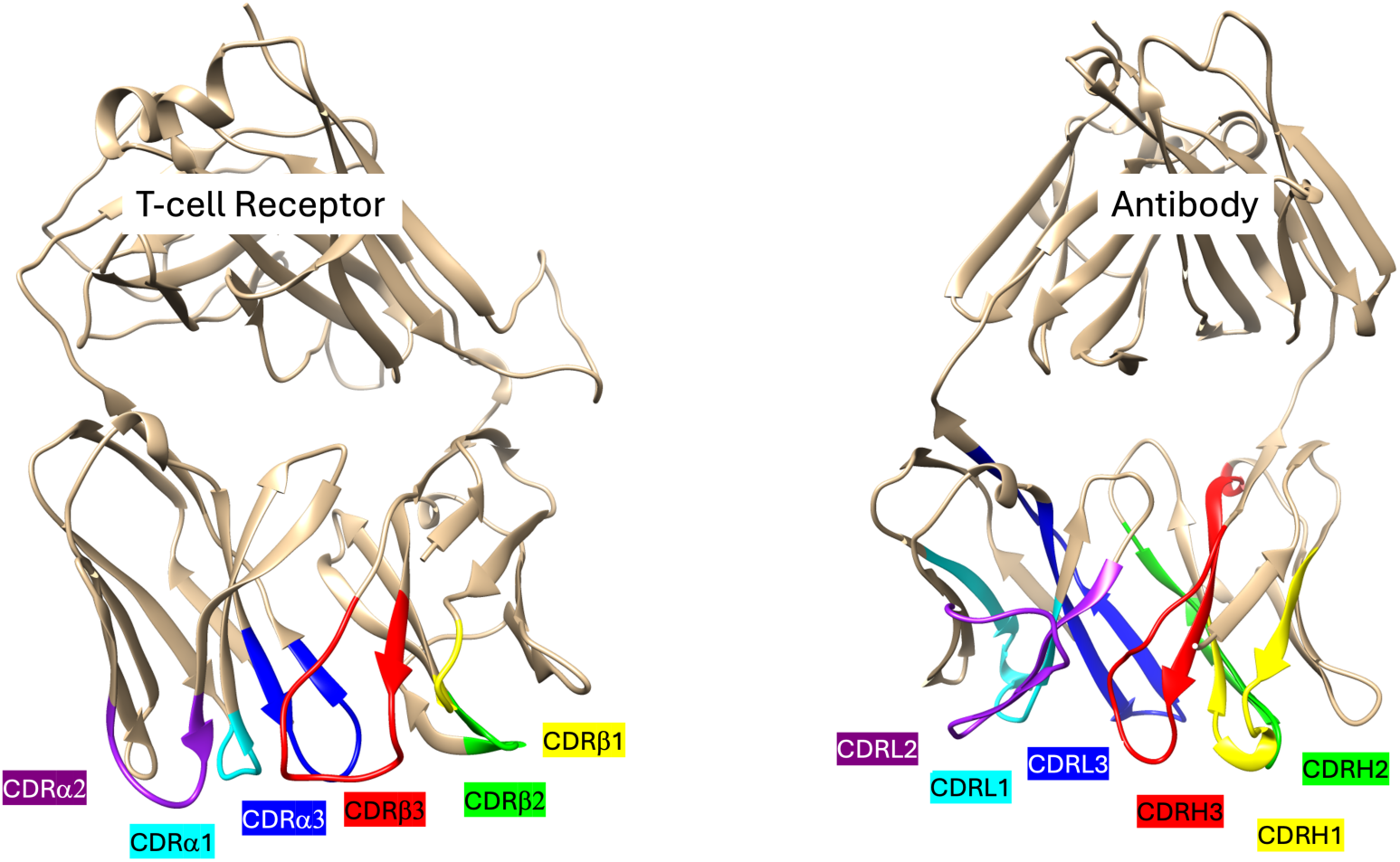
Structural parallels between TCRs (on the left) and antibodies (on the right). Analogous chains and CDRs are color-coded consistently.

Our systems of interest, TCRs and Abs, were selected because of their structural similarities, and the well-known challenges associated with accurately modeling their loops[5, 12, 13]. TCRs and Abs share a broadly similar 3D architecture (**Figure 3**): both are heterodimers containing six complementarity-determining regions (CDRs) that form their binding interfaces. As illustrated in **Figure 3**, CDRH1, CDRH2, and CDRH3 from the Ab heavy chain (H) correspond to CDRβ1, CDRβ2, and CDRβ3 of the TCR β chain; likewise, the three CDRs of the Ab light chain (L) align with those of the TCR α chain. Despite these structural parallels, Abs can bind to an extraordinarily wide range of molecular targets, including peptides, proteins, and haptens, whereas αβ TCRs recognize peptide antigens presented by MHC molecules.

RMSDs were calculated for the entire system as well as for the CDR loops, which are the most challenging regions and contain the key structural information.

As shown in **Figure 4**, AF3 workflow variants with MSA based on reduced sequence datasets performed comparably to the original AF3 for TCR modelling, including for the challenging CDR3β loop. On the contrary, MSA-free predictions performed markedly worse **(Figure 4a** and **Figure 4b)**, with an average RMSD considering all heavy atoms in the entire TCR of 10.5 ±6.6 Å, underscoring that some degree of evolutionary information remains essential for accurate TCR structure prediction (**Figure 4b**). Notably, UniRef90-TCR and UniRef-textmining based variant yields the lowest average RMSD for CDR3β among all tested configurations **(Figure 4)**, a particularly relevant result given that CDR3β is widely recognized as the most difficult TCR loop to model accurately[5, 13]. The results shown in **Figure 4** were generated using a single seed. Repeating the analysis with five different seeds produced consistent outcomes, with no statistically significant difference between results obtained from the original AF3 and the AF3 variants with reduced MSA, as shown in the **Supporting Information Fig. 1**. The RMSD differences across variants with reduced sequence datasets are not statistically significant (p >0.05, annotated as “ns” in **Figure 4a**), indicating that reduced sequence datasets for MSAs can effectively compete with the full AF3 reference database. Crucially, these reduced-MSA variants offer substantial computational advantages, enabling up to a 45-fold reduction in runtime. The AF3 variant with a reduced database of sequences, decreases the computation time from 15 minutes to less than 40 seconds (on a recent Dell Precision 3680 with 1 GPU NVIDIA RTX A4500 Ada Generation; 1 cpu used). RMSD values for each TCR in the validation set across all AF3 variants are provided in the **Supporting Information.**

**Figure 4.**
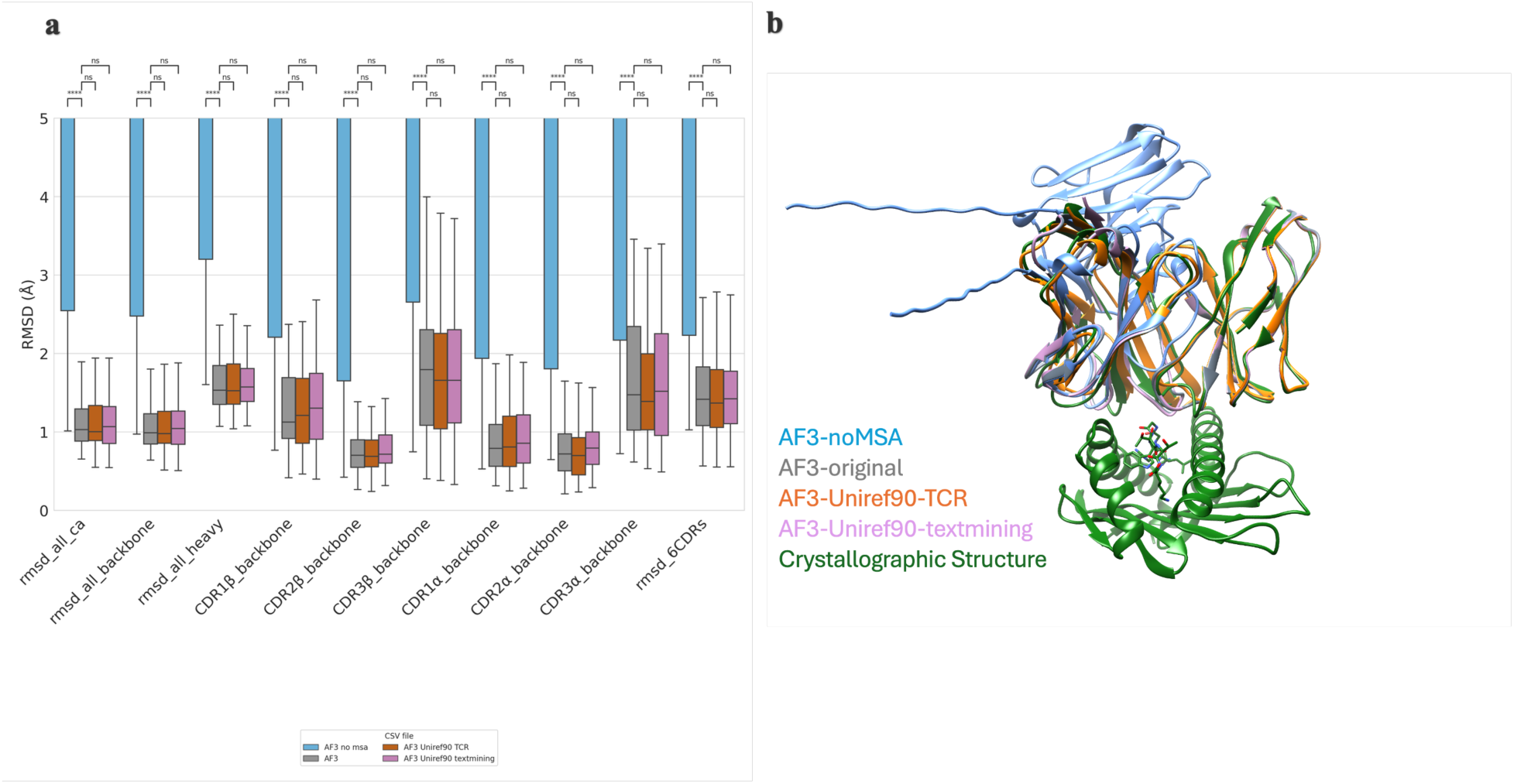
Analysis using different AF3 MSA variants for TCR modelling. **a)** RMSD analysis using one seed. We compared results of AF3 runs using four MSA configurations: no MSA (in blue), the default AF3 that uses among others UniRef90 (in gray), a UniRef90 subset selected according to AF3’s choices during TCR modelling from the VDJ database, i.e. UniRef-TCR (in orange), and a UniRef90 subset curated using TCR and antibody sequences, UniRef-textmining (in violet). The evaluation used a set of 77 TCRs for which just one TCR was accidentally used to also generate the sequence datasets for the MSA. RMSDs were calculated for the entire TCR receptor domain based on carbon alpha (rmsd_all_ca), backbone (rmsd_all_backbone) and all heavy atoms (rmsd_all_heavy). Backbone RMSD for the CDR1β (CDR1β_backbone), CDR2β (CDR2β_backbone), and CDR3β (CDR3β _backbone), CDR1a (CDR1a_backbone), CDR2a (CDR2a_backbone) and CDR3a (CDR3a _backbone). P-values calculated with Mann-Whitney-Wilcoxon test two-sided. The annotations used in the graph are ns: p>5.00e-02, *: 1.00e-02 < p <= 5.00e-02, **: 1.00e-03 < p <= 1.00e-02, ***: 1.00e-04 < p <= 1.00e-03 and ****: p <= 1.00e-04. **b)** Superimposition of the crystallographic structure (PDB code 7RRG) with structural models of the same TCR generated using four AF3-MSA variants: no MSA (in blue), AF3 (in gray), UniRef90-TCR (in orange), and UniRef90-textmining (in violet). Five seeds (and five inferences) were used, and the model with the best AF3 score was selected for analysis. The RMSD between the no-MSA variant model and the crystallographic structure is 13.5 Å. In this model, the long-coiled terminal regions are parts of the TCR variable domain not properly folded (and not to any signal peptide).

The correlation between TCR RMSDs obtained from the native and AF3 workflow variants can be seen in **Figure 5** and **Supporting Information Figures 2 and 3**. We observe that correlation between the RMSD obtained from AF3 (full databases) and AF3 workflow variant with UniRef-TCR database models is higher than 0.75 for all the different RMSD analysis including loop-specific RMSDs, indicating that expected prediction accuracy for each TCR is preserved regardless of whether the native or accelerated workflow is used, with both workflows reproducing the same overall error patterns and relative loop-specific modeling difficulties.

**Figure 5.**
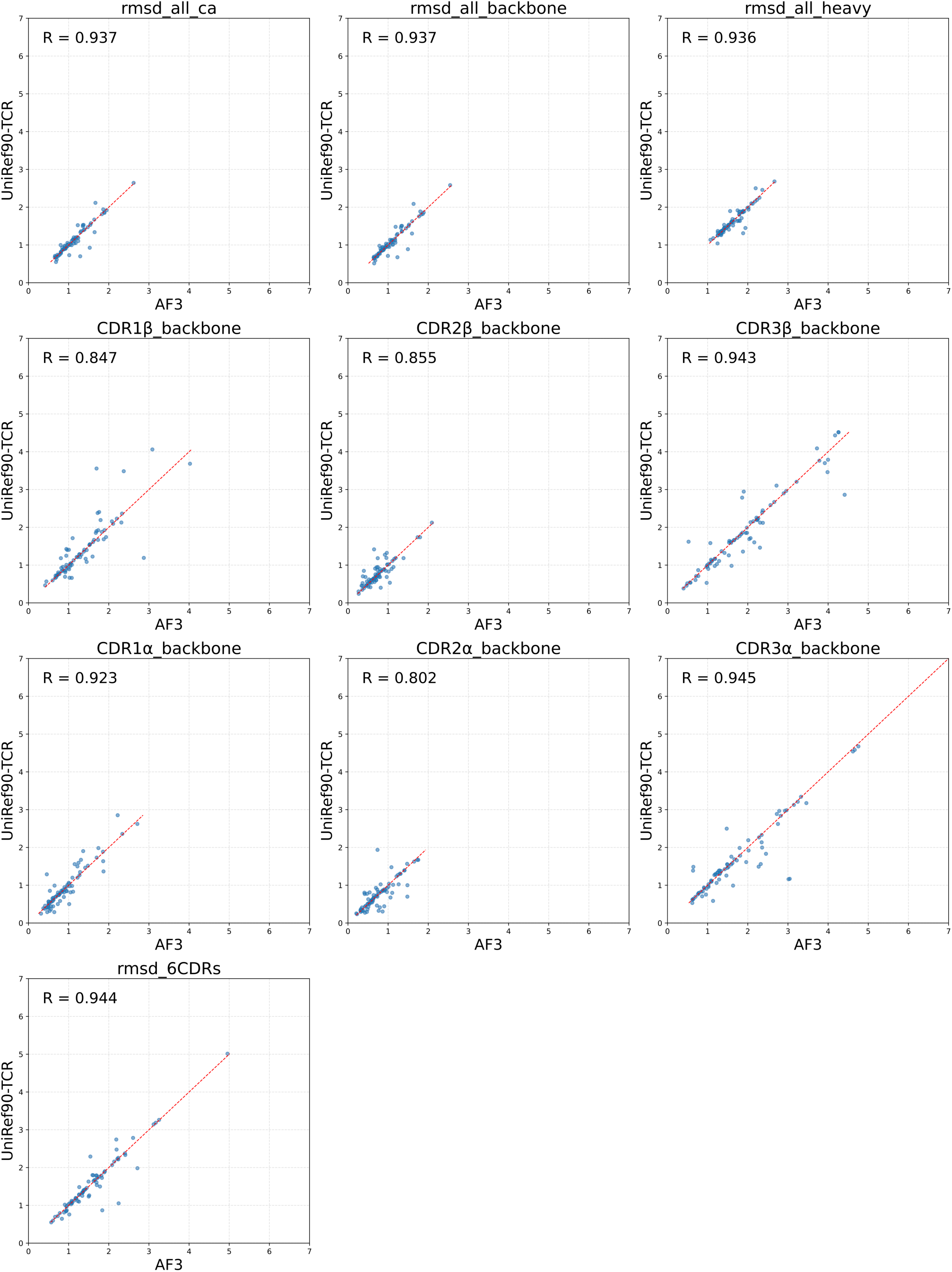
RMSD correlations for the native AF3 workflow and one variant using UniRef-TCR. RMSD were calculated between the experimental structure and the model generated by a given AF3 workflow, on the entire TCR based on heavy atoms (rmsd_all_heavy), on Cα atoms (rmsd_all_Ca), and on backbone atoms (rmsd_backbone), as well as for the backbone of the CDR1β (CDR1β _backbone), CDR2β (CDR1β _backbone), CDR3β (CDR1β _backbone), CDR1a (CDR1a_backbone), CDR2a (CDR2a_backbone), and CDR3a (CDR3a_backbone) loops.

The AF3 MSA pipeline was run either with its default databases or exclusively with the specialized databases on a set of 525 Abs from SAbDab (thus with an experimental 3D structure) that were not part of the AF3 training set. The inference was performed with 10 seeds. We then compared the RMSD of the backbone atoms of the models to their crystal structures. Whatever the native or accelerated workflows used, we observed a much higher RMSD for CDRH3 than any other CDR – the latter having a median RMSD below 1Å (**Figure 6a**). When comparing the RMSD for all CDRs between the models created with the accelerated (using the Uniref-Ab sequence dataset) and native AF3 workflows, the first showed marginally higher RMSD despite having a median RMSD with the same order of magnitude, indicating that the predictive performance of the two workflows is effectively indistinguishable. A high correlation could be observed between the 6 CDRs RMSD of the models generated with the native and this accelerated AF3 workflows (pearson correlation: 0.917) as well as between the native AF3 workflow and that using the AF3-textmining sequence dataset (pearson correlation: 0.871), **Figures 6b and c**, respectively. The correlation between CDRs’ RMSD obtained from the native and AF3 workflow variants can be seen in **Supporting Information Figures 5 and 6**. Thus, similarly to TCRs, the use of reduced-MSA protocols to model Ab structures reproduce the same overall error patterns and relative loop-specific modelling difficulties as the original AF3 workflow.

**Figure 6.**
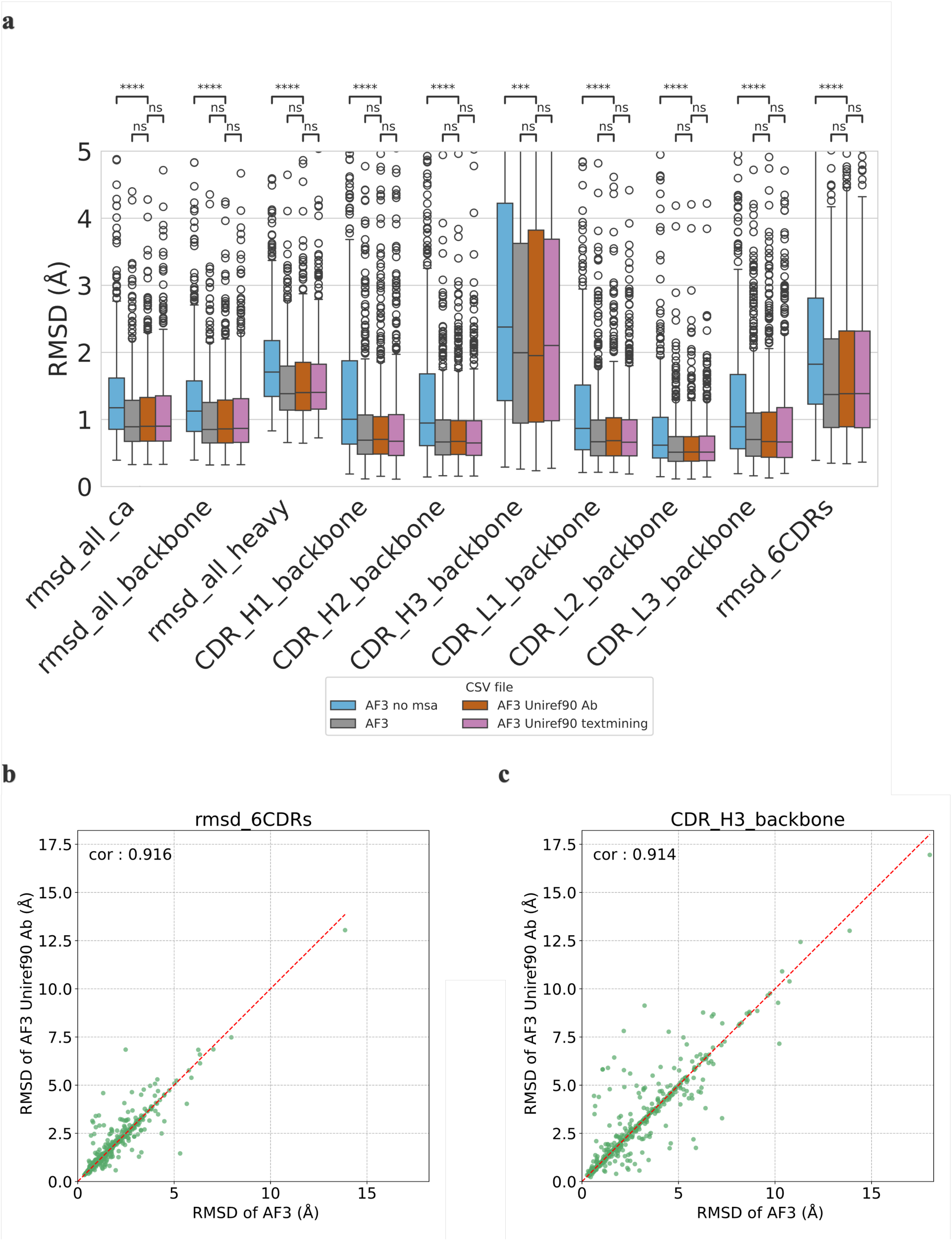
RMSD analysis using different AF3 workflow variants for Ab modelling. **a)** We compared results of AF3 runs using four MSA configurations: no MSA (in blue), the default AF3 that uses among others UniRef90 (in orange), a UniRef90 subset selected according to AF3’s MSA on Ab modelling of 2,490 structures, i.e. UniRef-Ab (in green), and a UniRef90 subset curated using TCR and antibody sequences, UniRef-textmining (in red). For evaluation, we used 525 Abs from SAbDab that were not included in AF3’s training set. RMSDs were calculated for the entire Ab’s Fv domain based on carbon alpha (rmsd_ca), backbone (rmsd_backbone), and all heavy atoms (rmsd_all heavy). Backbone RMSD for the CDR H1 (CDRH1_backbone), CDR H2 (CDRH2_backbone), and CDR H3 (CDRH3_backbone), CDR L1 (CDRL1_backbone), CDR L2 (CDRL2_backbone), and CDR L3 (CDRL3_backbone). The annotations used in the graph are ns: p > 5.00e-02, *: 1.00e-02 < p <= 5.00e-02, **: 1.00e-03 < p <= 1.00e-02, ***: 1.00e-04 < p <= 1.00e-03 and ****: p <= 1.00e-04. For a non-cropped Y axis see **Supporting Information** Figure 7. In **b)** and **c)** we observed a high correlation between the RMSD of all 6 CDRs between AF3 and Uniref-Ab or Uniref-textmining workflow variants.

### One seed is enough for high-accuracy modelling of TCR and Ab receptor domains

We analysed the relationship between the AF3 ranking scores - generated across all models, using 20 seeds and five inferences per seed applied to the TCR validation set – and the model accuracy (**Figure 7a-b**). We have observed some outliers, highlighted with an orange cross for which the RMSD using the 6CDRs was higher than 15 Angstrom and the ranking score was lower than 0.4. Surprisingly, for example, for TCR with PDB code 7N2Q and for seed 56970, the ranking scores range between 0.35 and 0.44 and the RMSD ranges between 0.7 and 16.1 Angstrom (**Figure 7c**). Our results showed that a high ranking score does not reliably correspond to an experimentally accurate prediction, nor does the top-ranked model necessarily represent the best-quality structure among those generated. Furthermore, increasing the number of seeds does not improve model accuracy, indicating that additional sampling offers no benefit in this context (Figure 7d).

**Figure 7.**
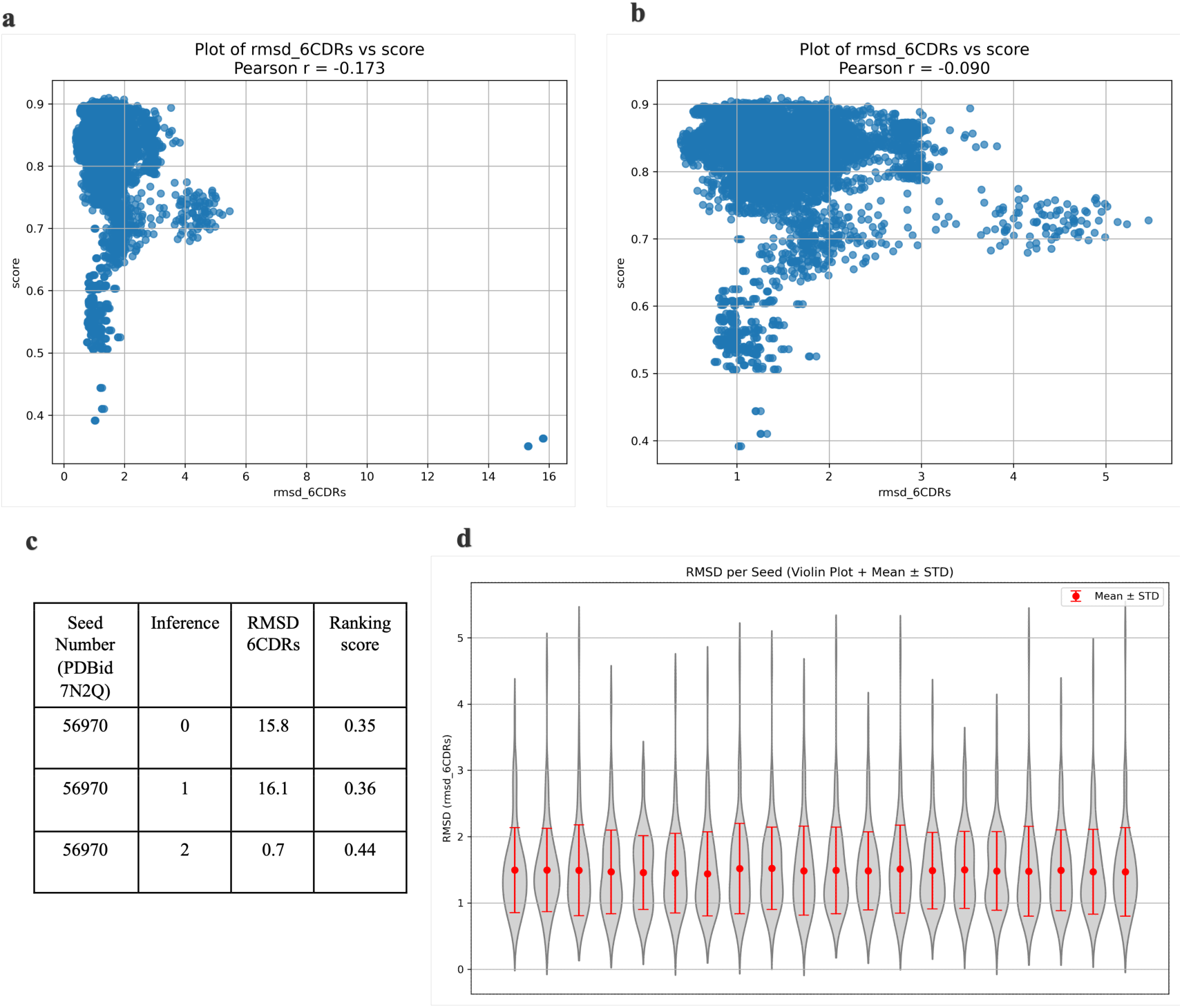
(**a-b**) Ranking score vs backbone RMSD calculated for the 6 CDR loops using UniRef-TCR sequence dataset. 20 seeds and 5 inferences per seed were used on a set of 77 TCR structures with known 3D that were not included in AF3 training. A) with outliers and B) without outliers. Ranking score = 0.8 × ipTM + 0.2 × pTM + 0.5 × disorder factor – 100 × has_clash as described in the code https://github.com/google-deepmind/alphafold3. **c)** Ranking score and RMSD for a PDB example: TCR 7n2q – 3 inferences for seed 56970. The ranking scores range between 0.35 and 0.44 and the RMSD for the 6CDRs between 0.7 and 16.1 Angstrom. **d)** RMSD of the 6 CDR loops as a function of the number of inference seeds used. Each run with n+1 seeds includes all seeds from the run with n seeds, plus one additional randomly selected seed. At each iteration, the RMSD is analysed relative to the best ranking score obtained across all preceding seeds and the newly added seed. RMSD analysis of TCR modelling using an AF3 variant with the UniRef90-TCR sequence dataset. A maximum of 20 random seeds were used.

We also analysed the AF3 ranking scores generated across all models, using 10 seeds and five inferences per seed for the Ab validation set (**Figure 8**). We found that for the same Ab, several models achieved low RMSD for CDRH3 or all CDRs but were not top ranked. So, suboptimal model selection (i.e. scoring failure) is observed. The AF3 ranking score showed poor correlation with CDR accuracy as measured by the RMSD, both for all CDRs (Pearson r = −0.297) and for CDRH3 (r = −0.247). Increasing the number of seeds to 20 did not improve CDR modelling or model selection, and the same trend was observed with the AF3 variants with reduced UniRef-Ab and UniRef-textmining sequence datasets (**Supplementary Figures**). These results indicate that the default AF3 ranking score, dominated by the ipTM term, inadequately reflects Ab model quality.

**Figure 8.**
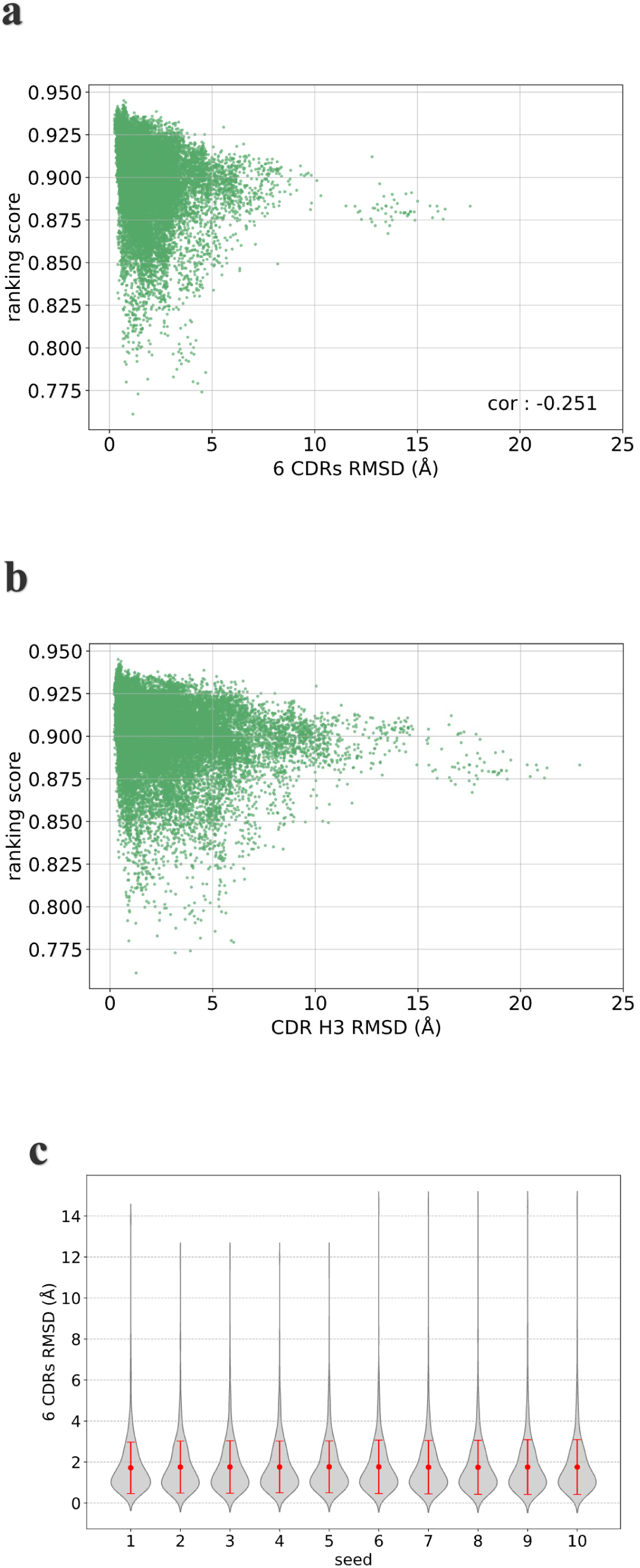
Analysis of the relationship between AF3’s ranking score and CDRs modelling accuracy. We assessed the correlation between the ranking score and **a**) the RMSD of all 6 CDRs and **b**) the RMSD of CDR H3. **c**) We plotted the evolution of the RMSD of all CDRs over the generation of models for 10 seeds, taking for each seed the best ranking score among the generated models up to that seed.

### Comparisons with faster-than-AF3 state-of-the-art method shows that AF3 using a reduced MSA achieves comparable or superior accuracy while offering superior speed

We analyzed the accuracy of AF3 and AF3 variants with reduced MSA modelling TCR against competitors (**Figure 9 a-b**). The two state-of-the-art competitors used were Boltz2 and TCRBuilder2. Boltz2, similarly to AF3, is a general-purpose biomolecular structure predictor, and TCRBuilder2 is a specialized, computationally efficient tool optimized specifically for TCR structure prediction. We compared AF3 with Boltz2 in **Figure 9a)** and AF3 with TCRBuilder2 in **Figure 9b)**, as the sizes of the evaluation sets differ. Because Boltz2 was published later, a smaller dataset was used for this comparison to ensure that none of the evaluated structures were included in the training data of any software (see **Methods**).

**Figure 9.**
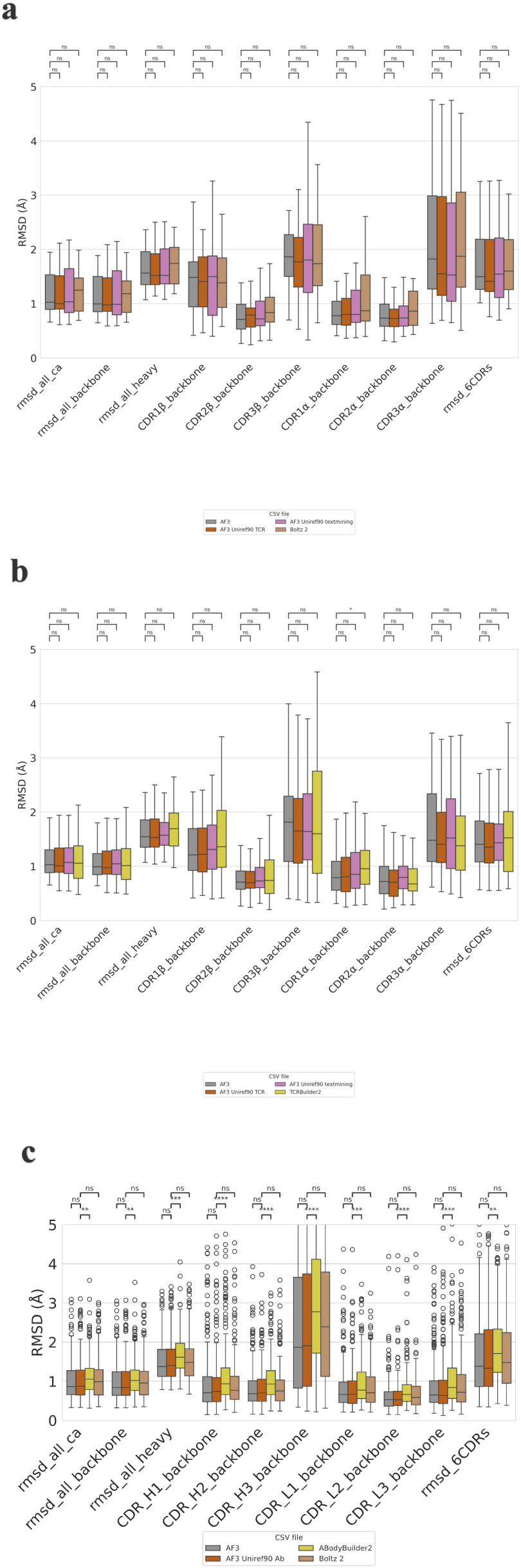
RMSD analysis using different AF3 MSA variants for TCR modelling and one seed, and comparison with state-of-the-art approaches. We compared results of AF3 runs using three MSA configurations: the default AF3 that uses among others UniRef90 (in gray), a UniRef90 subset selected according to AF3’s choices during TCR modelling from the VDJ database, i.e. UniRef-TCR (in orange), and a UniRef90 subset curated using TCR and antibody sequences, UniRef-textmining (in violet). Comparison with Boltz2 in **a)** and with TCRBuilder2 in **b).** RMSDs were calculated for the entire TCR receptor domain based on carbon alpha (rmsd_all_ca), backbone (rmsd_all_backbone) and all heavy atoms (rmsd_all_heavy). Backbone RMSD for the CDR1β (CDR1β_backbone), CDR2β (CDR2β_backbone), and CDR3β (CDR3β _backbone), CDR1a (CDR1a_backbone), CDR2a (CDR2a_backbone) and CDR3a (CDR3a _backbone). P-values calculated with Mann-Whitney-Wilcoxon test two-sided. **c)** RMSD was computed on the Fv domain of 236 Abs and compared to AF3 UniRef90 Ab, on carbon alpha (rmsd_ca), backbone (rmsd_backbone), and all heavy atoms (rmsd_all heavy). Backbone RMSD for the CDR H1 (CDRH1_backbone), CDR H2 (CDRH2_backbone), and CDR H3 (CDRH3_backbone), CDR L1 (CDRL1_backbone), CDR L2 (CDRL2_backbone), and CDR L3 (CDRL3_backbone) were also computed. The annotations used in the graph are ns: p <= 1.00e+00, *: 1.00e-02 < p <= 5.00e-02, **: 1.00e-03 < p <= 1.00e-02, ***: 1.00e-04 < p <= 1.00e-03 and ****: p <= 1.00e-04.

We observe that all approaches show similar accuracy when modelling TCRs. All p-values are non-significant, except for CDR1α, where AF3 performs significantly better than TCRBuilder2 (**Figure 9b**). In terms of computational efficiency, AF3 variants with reduced databases, Boltz-2 and TCRBuilder2 are faster than AlphaFold 3. Predictions are typically completed in seconds (TCR Builder 2 and AF3 variants with reduced databases) to a few minutes (Boltz-2), whereas for AF3 generally requires several minutes and higher computational resources.

The same anaylsis of competitors was also performed for Abs, as mentioned previously, to have a validation set that doesn’t overlap with the training set of Boltz2 a smaller validation set was selected. We observed that ABodyBuilder 2 was performing significantly worse than AF3. While Boltz 2 had also less accurate prediction than AF3, it was only significant for CDR L1 (**Figure 9c**).

### AF3 parameters optimization during inference allows for substantial speed up for modelling small proteins

First, by reducing the default GPU memory pre-allocation to 10% instead of 95% we were able to run up to 9 inferences in parallel on a single 24Gb GPU. While these numbers are specific to the hardware used (10% allowed us to generate models of up to 312 tokens), we hypothesize that on a GPU with more dedicated memory we could run even more inferences in parallel. Parallelizing inferences allows to reduce the featurization and the first seed inference time, respectively from 7 to 1 second and from 36 to 23 seconds (**Figure 10a**), but not subsequent seeds. This probably means that only the model compilation and featurization which are handled by the CPU were accelerated through this parallelization.

**Figure 10.**
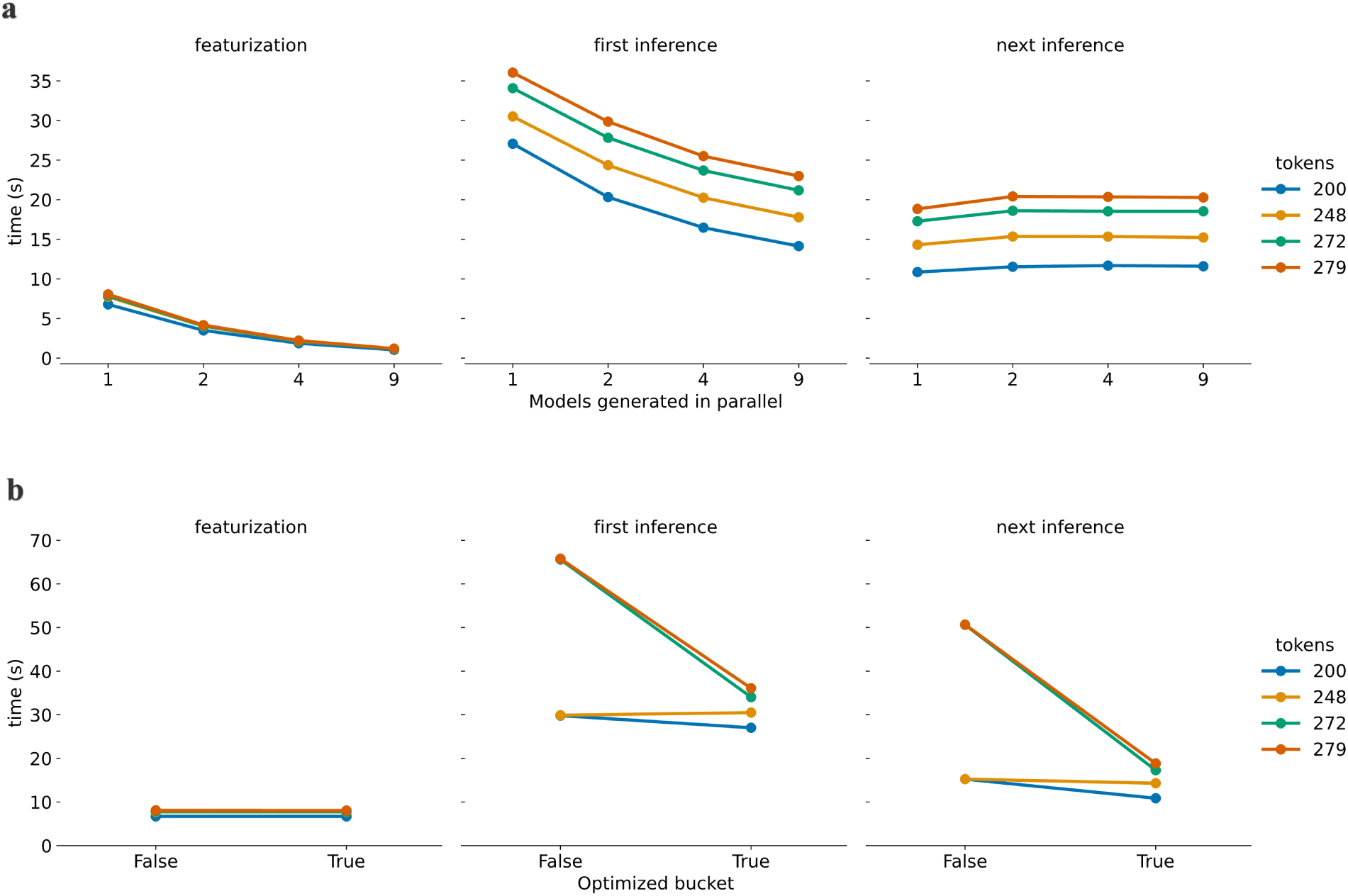
Temporal analysis of GPU optimized AF3 inference for 2 seeds with different Ab length in tokens (1 token equals 1 amino acid). We compare relative speed for different input lengths under the same conditions for parallelization with optimized buckets **a)** or only for bucket optimization **b)**.

To further reduce inference time, we tailored bucket size to the exact length of our inputs (1 amino acid = 1 token) and observed a speed up proportional to the amount of padding tokens eliminated (**Figure 10 b**). The most striking example was a 272 tokens Ab that would by default use a bucket of 512 tokens resulting in 240 padding tokens. The inference of the first and second seeds respectively took 65.5 and 50.5 seconds with default bucket while it only took 34 and 17.3 seconds with optimized bucket size, this showed a roughly flat 30 seconds time gain per seed inference. This optimization is not hardware specific and would provide significant speed up to any processus needing to generate large number of small models with AF3. We did not analyse more seeds because all subsequent seeds after the second one are identical to it (**Supporting Information Figure 8**).

## Discussion

Despite the reduced influence of the MSA on model construction in AF3, the MSA component remains the slowest step in the workflow, representing more than 90% of the end-to-end execution time. Consequently, MSA generation imposes a disproportionate runtime and emerges as a bottleneck in AF3.

To largely decrease the end to end execution time of AF3 to generate structural models of TCR or Ab receptor domains, four different AF3 MSA variants were developed: no MSA, the original AF3 setup with the full UniRef90 database, a UniRef90 subset selected according to AF3’s choices during TCR and Ab modelling (UniRef90-TCR and UniRef90-Ab, respectively) and a non-exhaustive UniRef90 subset curated using TCR and Ab sequences (UniRef90-textmining).

All the models obtained with the AF3 MSA-free variant are significantly worse than the ones obtained with the reduced databases and original AF3 (p-value < 0.0001) for the modelling of TCRs, indicating that MSA is needed for the modelling of the TCRs. Conversely, we observed that for Ab modelling the MSA-free variant can be used while keeping accuracy near experimental resolution (except for about 11 structures, 6 CDRs RMSD > 10Å, see **Supporting Information**) – at the cost of a very marginal yet significant loss in accuracy compared to the native AF3 workflow. We hypothesize that these differences arise partly because far fewer TCR experimental 3D structures were available for AF3 training compared to antibodies. Although some TCRs are better modelled than others in the MSA-free variant, we do not observe that the ones with low RMSD have better representation of their genes in the 3D structures that were used for AF3 training (data not shown).

Despite using highly size-reduced subsets of UniRef90 for MSA, all our accelerated Af3 workflows lead to results comparable to the native AF3 workflow. The RMSD differences are comparable indicating that reduced-database MSAs can effectively compete with the full AF3 reference database for some conserved families/domains of proteins, including TCRs and Abs.

We observed that the Pearson correlation between the original AF3 model and each reduced-MSA AF3 variant for TCR and Ab modelling exceeds 0.75 across all RMSD evaluations, including the per-loop analyses. In addition to reflecting methodological consistency, a high Pearson correlation suggests three important points. First, the structural variability captured by the models is largely driven by intrinsic features of the TCRs and Abs, such as loop length, conformational flexibility, or limited evolutionary signal, rather than by the specific size of the MSA database used. Second, the strong correlation implies that reducing the MSA complexity does not introduce new biases or failure modes; the variants remain aligned with AF3’s ranking of “easy” versus “difficult” cases to model. Third, this stability supports the interpretability and robustness of the reduced-database approach, reinforcing that computational efficiency can be gained without altering the fundamental behaviour of the prediction algorithm.

Crucially, these reduced-MSA variants offer substantial computational advantages, enabling up to a 45-fold reduction in runtime. AF3 variant with a reduced database of sequences, decreases the computation time from 17 minutes (11 minutes for Abs) to less than 40 seconds (10 seconds for Abs, with 8 cpus used) (Machine Dell Precision 3680 with 1 GPU NVIDIA RTX 4500 Ada Generation, 1 cpu used).

Our results showed that the AF3 ranking score does not correlate well with loop RMSD for either TCRs or Abs. We therefore focused our analysis on loop regions, which represent the most challenging to model in these structures, whereas framework regions are highly conserved and typically achieve RMSDs of ∼1 Å. Across different numbers of seeds and inference runs, we consistently observed that the top-ranked models were not those with the lowest loop RMSD, leading to suboptimal model selection (i.e. scoring failure). Importantly, increasing the number of seeds did not improve the average RMSD, indicating that generating models for more than one seed provides little to no benefit. In this context, AF3 with a reduced MSA and a single seed represents an excellent trade-off between computational efficiency and accuracy for TCR and Ab modeling.

While MSA information is still used in AF3, its contribution appears substantially reduced compared with AF2, suggesting additional opportunities for optimizing the MSA stage without sacrificing accuracy. Although several faster competing approaches exist and have demonstrated strong performance in their original publications, they have already been extensively compared to AF3 by their developers. AF3 relies on JackHMMER to generate MSAs, which is computationally expensive but produces deep, high-quality alignments. Faster alternatives, such as MMseqs2—used in the ColabFold API based on AF2—could further reduce runtime. However, the use of such a tool puts an extra layer of complexity on the user’s workflow, while our method is simple and merely requires the user to use the dedicated AF3 options for specific databases and to create a specialized database of sequences.

We benchmarked AF3 variants against other state-of-the-art modelling tools to compare their predictive accuracy. We observe that TCRBuilder2, Boltz-2, AF3 and AF3 variants show similar accuracy when modelling TCRs. Differences in model accuracy are non-significant, except for CDR1α, where AF3 performs significantly better than TCRBuilder2. For modelling Ab, AF3 performed better than AbodyBuilder2 and similar to Boltz2. Boltz-2 was competitive in terms of accuracy but significantly more time consuming than our AF3 variants, making the latter a good choice for TCR and Ab modelling. Eventually, we also found some AF3 specific parameters to reduce the inference time down to 20 seconds for Abs.

In conclusion, we analyzed AF3 performance for TCR and Ab modeling, varying the MSA generation and inference phase. Using reduced subset of sequences from UniRef90 greatly speeds up AF3, achieving ∼45× faster runtimes on a desktop while maintaining near-experimental accuracy, enabling scalable structural immunology.

## Methods

### TCR and Antibody set used for MSA analysis

A non-redundant set of 3,213 TCR sequences, derived from the VDJ database[10, 11], was used to identify AF3-selected sequences for multiple sequence alignment (MSA), covering specificities for 200 different antigens. Detailed information on all TCRs, including gene composition and peptide specificity, is provided in Suplementary Table 9 of Mayol-Rullan *et al[7]*.

The SAbDab [14, 15] website was used to retrieve a non-redundant set of Abs with at most 95% sequence identity with paired chains (Heavy and Light) on the 20^th^ of January 2026. All bound and unbound structures between 1^st^ of January 2022 and the 15^th^ of October 2025 were removed from this set as well as the structures with the same concatenated CDR loops as structures between 1^st^ of January 2022 and the 15^th^ of October 2025. This resulted in a set of 2,490 diverse Ab sequences.

### Custom Uniref90-derived databases creation

Multiple sequence alignment (MSA) of the TCR sequences was performed using JackHMMER [9] (EVALUE=1e-3, NUM_ITER=3) against the UniRef90 database[8, 17]. All unique aligned sequences were aggregated into a new database called **UniRef-TCR** composed of 261’780 sequences, among which 261’055 were aligned with the alpha chain and 208’047 with the beta chain. The EVALUE threshold of 1e-3 used here is less stringent than the 1e-4 EVALUE typically used in AF3 MSA runs. This relaxed criterion allows for faster sequence expansion and the identification of more distant homologs. The goal is to avoid overly constraining the sequence space, providing AF with a reduced set but still with a broader diversity of sequences.

Another subset of UniRef90 was created by searching in the database for the words “t-cell receptor”, “t cell receptor”, “Tcell receptor”, “immunoglobulin”, “antibody”, “nanobody” in the sequences header. This resulted in a non-exhaustive set of sequences with a total of 41’055 sequences gathered this way. We called this new database **UniRef-textmining.**

### Validation dataset

TCR structures bound to a pMHC class I uploaded to the Protein Data Bank (PDB) between the 1st of January 2022 and 12 November 2025 (outside of AlphaFold3 training set) were collected. As our objective is to model TCR receptors alone to predict their specificity just TCR-pMHC class I structures were considered and 77 structures were retained. The PDB codes of these structures are 8rj5,9d95,9gv6,9gv7,9iky,9j4t,9j4u,9k2i,9k2r,9ru5,9rup,9rxm,8en8,8enh,8eo8,8qfy,8rlt,8rlu,8rlv,8rym,8r yn,8ryo,8ryp,8ryq,8v4z,8v50,8wte,8wul,8ye4,9c3e,7q99,7q9a,7q9b,8dnt,8es9,8i5c,8shi,6zkw,6zkx,6zky, 6zkz,7dzm,7dzn,7n2n,7n2o,7n2p,7n2q,7n2r,7n2s,7n4k,7n5c,7n5p,7na5,7ndq,7ndt,7ndu,7nme,7nmf,7nm g,7ow5,7ow6,7pb2,7pbc,7pbe,7pdw,7phr,7qpj,7r80,7rk7,7rm4,7rrg,7su9,8cx4,8d5q,8gvb,8gvg,8gvi.

Ab structures bound to a peptide or protein antigen according to SAbDab and added between the 1st of January 2022 to 15th of October 2025 were collected. Structures with a resolution above 3Å and or an R-free value superior to 0.3 were excluded. Only structures with two different chains as their heavy and light chains were kept. Moreover, only the structures without missing residues were analysed resulting in a total of 525 structures. The PDB code for the Ab structures are 9r2e,9fyt,7yzj,8tft,7trh,8zpq,8vvk,7s5p,8v13,8sdh,9fik,7ox1,7udh,8tq9,8xx0,8tfl,8zd5,9cds,7zoz,8qxc,7w sl,8uky,8s6t,7y0g,9ghj,8rtw,7qt2,7qt0,8d1t,9axn,9axo,7qt3,8xk6,8g2o,8qya,8z39,8z2v,8z3a,7t0l,8yxi,8tq 5,8ipc,7ufn,7ufq,7ufo,9h4s,7ue9,9iut,9i1q,8f0l,8h8q,8dk6,8dcn,7t0f,8sip,8p6i,8s6v,9cpx,7zxk,9cpu,9cpv, 8cwj,8b7h,8rmx,7uvs,8z95,8up2,9ggp,9uvi,7yru,8hrd,7u5b,8vy4,9bqw,7t7b,8e1g,8bse,8qrg,9oas,7xsc,8d 9z,7s5r,7ycl,8tbq,9jqv,9ml9,8f60,8f6o,8zxw,8f6l,8u03,8u08,8wsp,8tzw,7ox3,8yor,7yds,8yhz,8og0,8u31, 8u32,9j8a,9ml8,8w86,8ruu,8c3v,9c7d,8cwi,8cz5,8vtd,7uij,8urf,8s73,7uzc,8jyr,9gwt,8gho,7wvl,8jyq,9npi, 9bdi,8sjk,8acf,9bdh,8gy5,8x0y,7u0a,7qu1,8bbo,8dao,8gb6,7xjf,8tqa,7yk4,8too,8tuz,9blx,8ffe,7tow,9lrf,7 x63,7xik,8v5q,8byu,8fhy,8g8a,7sl5,7xy8,8yx1,7wnb,7st5,8zu9,7zf8,7wn2,8elq,9c1s,9fm0,7wbz,9bns,8hn 6,8eli,7xeg,8ulj,9c7f,8f9t,8dxt,7tn0,8thn,8sj4,8fb5,8fa7,7y0c,8txp,9dx6,7y3o,8fb8,8fb7,8bk2,7urq,7fjc,9d ss,9dst,8fa9,8f9v,9cr9,7uxl,8f9u,8fan,8rp8,7yv1,7zqt,9jcy,9pwn,7yxu,9njy,7wn8,8f5i,7z2m,8fba,8d6z,7s sc,9f91,8g8c,8dtx,7unb,7t86,8fdc,7xrz,9avo,8dn6,8jnk,9cb5,8dn7,8vc7,9dl0,8caf,9mxi,8d29,8vm8,9d41, 8slb,9dbo,8ux6,8wsn,7skz,9gox,9goy,9dru,9oaq,8w0v,7uvi,8d47,9oao,8bh5,9byf,9rm2,8qh0,7s7i,7t25,8z 2e,8b9v,7str,7s5q,7ul1,8cbf,8w0w,9pix,8hn7,7z3w,8tui,8tv3,8gf2,7tp3,8dtt,8q3j,9bjg,8fax,8a44,8fdd,8t9 z,9bjh,8gb8,8dfg,9au1,7x2h,8be1,9cqb,8us8,7wlw,7zwi,7tuf,7y3j,8v2e,8ts0,7uzd,9mer,9dq4,7u64,7sa6,9 g6r,8rmo,8ja5,8eb2,8ds7,9c7x,8szy,7r58,9bt5,8bg1,8ee0,8ib1,9b2w,8che,7uen,8fat,8yx9,7txw,7wkx,7ue m,8fas,7uqc,8bbh,7ucx,8t59,8t51,7u63,7u8c,7u62,9je3,7vaz,8b50,8fg0,9lf8,9dh2,8tbb,7uvh,7uvf,7t87,8g 2m,8g8n,8gye,7ut3,9f90,8rwb,7wg3,8oni,8ds5,9g6s,9jbq,7tcq,7quh,9q8l,7u0e,9cqa,7x8p,8x0t,7x8t,9nph, 9mqv,7u09,8sga,8ykt,8uzp,8s72,8r4n,8bsf,8tmz,8dgu,8av9,8sdf,8fbw,7sts,8dgw,7y0v,7sx7,8d36,8vdl,8tq 7,9lra,8hpk,7ycn,9c6y,8zyf,8siq,7u2d,7u2e,8ahn,7tr4,7tp4,8sgn,8rpb,9atm,8s6m,7z0y,7wph,8dcc,7ul0,7z 0x,8el2,8d3a,7u8e,8udz,9jd1,8eoo,7ttm,8oxw,9fys,8vvl,7fgk,7myt,7ra7,7rpt,7s08,7s0j,7sjm,7tqa,7tuh,7tu s,7u0b,7u0k,7u5g,7u61,7uce,7ued,7uyl,7v3q,7ven,7vmz,7wvi,7zf6,7zff,8awl,8bf0,8bjz,8c67,8c7v,8cbx,8 d01,8d54,8dkf,8doz,8eez,8ef1,8ef2,8ef3,8ek6,8eq6,8eqa,8eqc,8f2t,8fwf,8fxj,8fzo,8fzp,8g19,8g1b,8g1c,8 gbv,8gbw,8gbx,8gc0,8gdo,8gkk,8gsi,8gv1,8hyl,8iqp,8iqs,8jbj,8jep,8jeq,8oju,8p2t,8p88,8ppm,8pul,8rel,8 rpe,8rtx,8ry2,8s6e,8sef,8sgg,8shv,8sve,8t0o,8t8i,8tca,8tfe,8ti4,8tjt,8tn4,8tv2,8uhh,8uhs,8uio,8uki,8ulh,8v 4i,8vbq,8vbr,8veg,8vfu,8vk2,8vlt,8vs8,8vtp,8vvb,8w9h,8y2k,8ywp,8z4c,8zay,9axq,9axs,9b76,9b88,9b8g,9c9k,9dmf,9ds3,9dsc,9E+69,9eci,9ecx,9ecy,9f18,9fqo,9gff,9gfg,9gfh,9gfl,9gjv,9gp2,9ii9,9iy0,9luk,9mi7,9mjc,9mji,9mk4,9n6u,9n7k,9n89,9pyf.

### AlphaFold3 model generation with custom databases

Structures modelled with AlphaFold3, used 5 inferences per seed and the 20 following randomly chosen seeds: 9986, 67719, 29960, 58505, 56970, 55176, 90764, 44813, 71819, 36501, 58009, 18970, 90779, 1948, 91807, 65829, 92455, 87079, 11815, 39719. When custom sequence databases were used, if a AF3 original databases needed to be removed, only the first two lines (the sequence header and the first line of the sequence) were kept as AlphaFold3 would throw an error when trying to use empty files as databases. The removed databases were bfd-first_non_consensus_sequences.fasta, mgy_clusters_2022_05.fa and uniprot_all_2021_04.fa.

The UniRef-TCR, UniRef-Ab selection and UniRef-textmining custom databases were temporarily renamed as the larger database they were issued from: uniref90_2022_05.fa when producing MSA through AF3. AF3 MSA pipeline and inference were run independently for optimization purposes. To accelerate model generation, the slurm queueing system was used to run batches of 4 MSA pipeline in parallel using 6 cpus each. Inference was run on a single Nvidia RTX 4500 Ada Generation GPU. Generating both chains of anantigen binding Fragment (Fab) of an antibody took on average 30 seconds for a single seed and 170 seconds for 10 seeds. Generating both chains of TCR took on average 40 seconds for a single seed.

### Root Mean Square Deviation (RMSD) calculation

To superimpose and compute the RMSD of TCRs and Ab models, the Biopython [18] package was used. Any crystal structures with missing residues were removed from the RMSD analysis. For a more accurate and consistent comparison, models were superimposed on their framework residues (thus excluding the structurally variable CDR loops) rather than on the whole structure and RMSD was computed either on single Complementary Determining Region’s (CDR) residues, on a set of all CDRs residues or on all residues of the model. To determine CDRs and framework residues the ANARCI software was used[16]. For TCRs, the IMGT numbering scheme was used while the Chothia numbering scheme was preferred for antibodies.

### Analysis of CDRs RMSD across multiple seeds

To compare the evolution of the RMSD of CDRs across multiple seeds we imitated AlphaFold3 behaviour by taking the model with the highest ranking score for each seeds taken into consideration (e.g., after 4 seeds, the highest score among the 20 inferences). RMSD of all CDRs residues were then computed. For Abs, only 10 seeds were generated.

### Comparison of AF3 with TCRBuilder2 and Boltz2

TCRBuilder2 was applied to model AF3 with default parameters and the AF3 validation set described above. Boltz2 was applied to model a subset of TCRs from the validation set. Structures published between 1^st^ January 2022 and July 2025 were excluded from the original validation set because Boltz2 was released after AF3, and some of the PDB structures used to validate AF3 were already included in Boltz2’s training data. Boltz2 was applied with 10 recycling steps and 25 diffusion samples as recommended for comparison with AF3 in the Boltz2 GitHub repository https://github.com/jwohlwend/boltz/blob/main/docs/prediction.md.

### AF3 inference optimization (GPU optimization)

Bucket optimization comparison was performed with the --buckets argument set to the exact same number of tokens as the input or left to the default bucket sizes (256 or 512 tokens). Parallelization was performed with optimized buckets, by creating an AlphaFold docker image with 10% pre-allocated memory (Dockerfile : ENV XLA_CLIENT_MEM_FRACTION=0.10) then inferences were launched in parallel. To assess the overall speed up a set of 100 random Abs from the validation set were modelled with optimized parameters. Featurization and seed inference times were retrieved through the logs of AF3. Note that computing 9 inferences in parallel necessitated about 40Go of RAM usage.

