## Supplemental Document for "Rapid and Reliable Structural Modeling of Adaptive Immune Receptors Using an Optimized AlphaFold3 workflow"

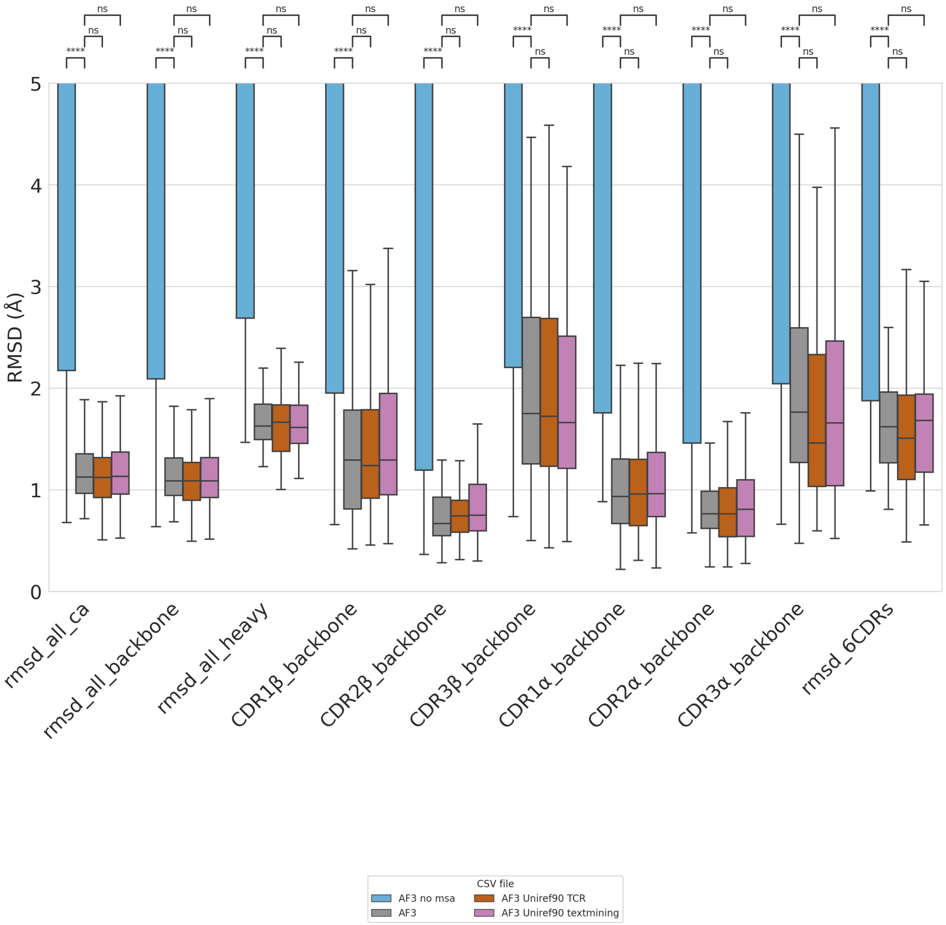


**SI Figure 1**. RMSD analysis using different AF3 MSA variants for TCR modelling and five seeds. We compared results of AF3 runs using four MSA configurations: no MSA (in blue), the default AF3 that uses among others UniRef90 (in gray), a UniRef90 subset selected according to AF3's choices during TCR modelling from the VDJ database, i.e. UniRef-TCR (in orange), and a UniRef90 subset curated using TCR and antibody sequences, UniRef-textmining (in violet). The evaluation used a set of 77 TCRs for which just one TCR was accidentally used to also generate the sequence datasets for the MSA. RMSDs were calculated for the entire TCR receptor domain based on carbon alpha (rmsd_all_ca), backbone (rmsd_all_backbone) and all heavy atoms (rmsd_all_heavy). Backbone RMSD for the CDR1β (CDR1β_backbone), CDR2β (CDR2β_backbone), and CDR3β (CDR3β _backbone), CDR1a (CDR1a_backbone), CDR2a (CDR2a_backbone) and CDR3a (CDR3a _backbone). P-values calculated with Mann-Whitney-Wilcoxon test two-sided. The annotations used in the graph are ns: p <= 1.00e+00, *: 1.00e-02 < p <= 5.00e-02, **: 1.00e-03 < p <= 1.00e-02, ***: 1.00e-04 < p <= 1.00e-03 and ****: p <= 1.00e-04.

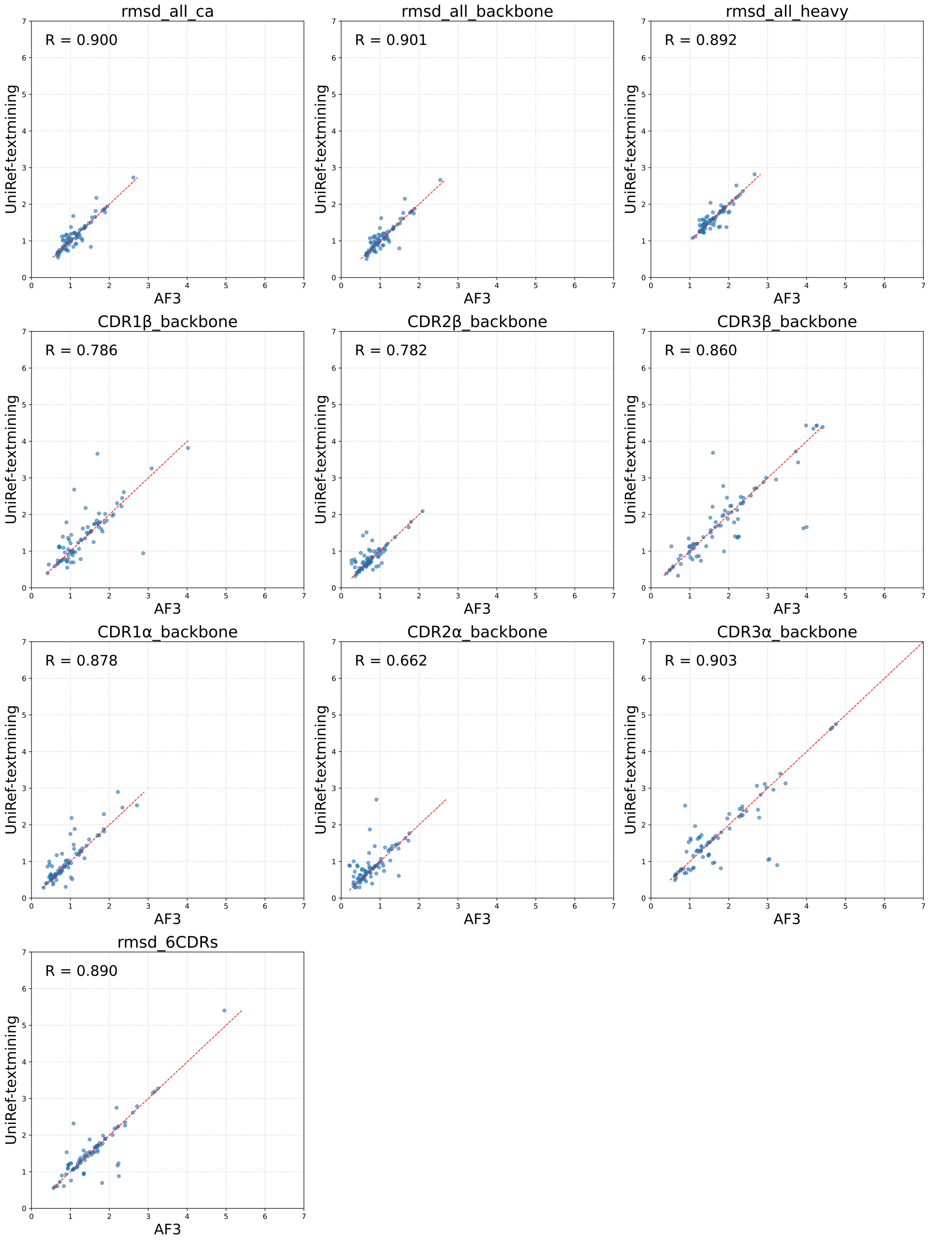


**SI Figure 2.** RMSD correlations for TCR modelling using the native AF3 workflow and one variant using **UniRef-textmining**. RMSD were calculated between the experimental structure and the model generated by a given AF3 workflow, on the entire TCR based on heavy atoms (rmsd_all_heavy), on Cα atoms (rmsd_all_Ca), and on backbone atoms (rmsd_backbone), as well as for the backbone of the CDR1β (CDR1β _backbone), CDR2β (CDR1β _backbone), CDR3β (CDR1β _backbone), CDR1a (CDR1a_backbone), CDR2a (CDR2a_backbone), and CDR3a (CDR3a_backbone) loops.

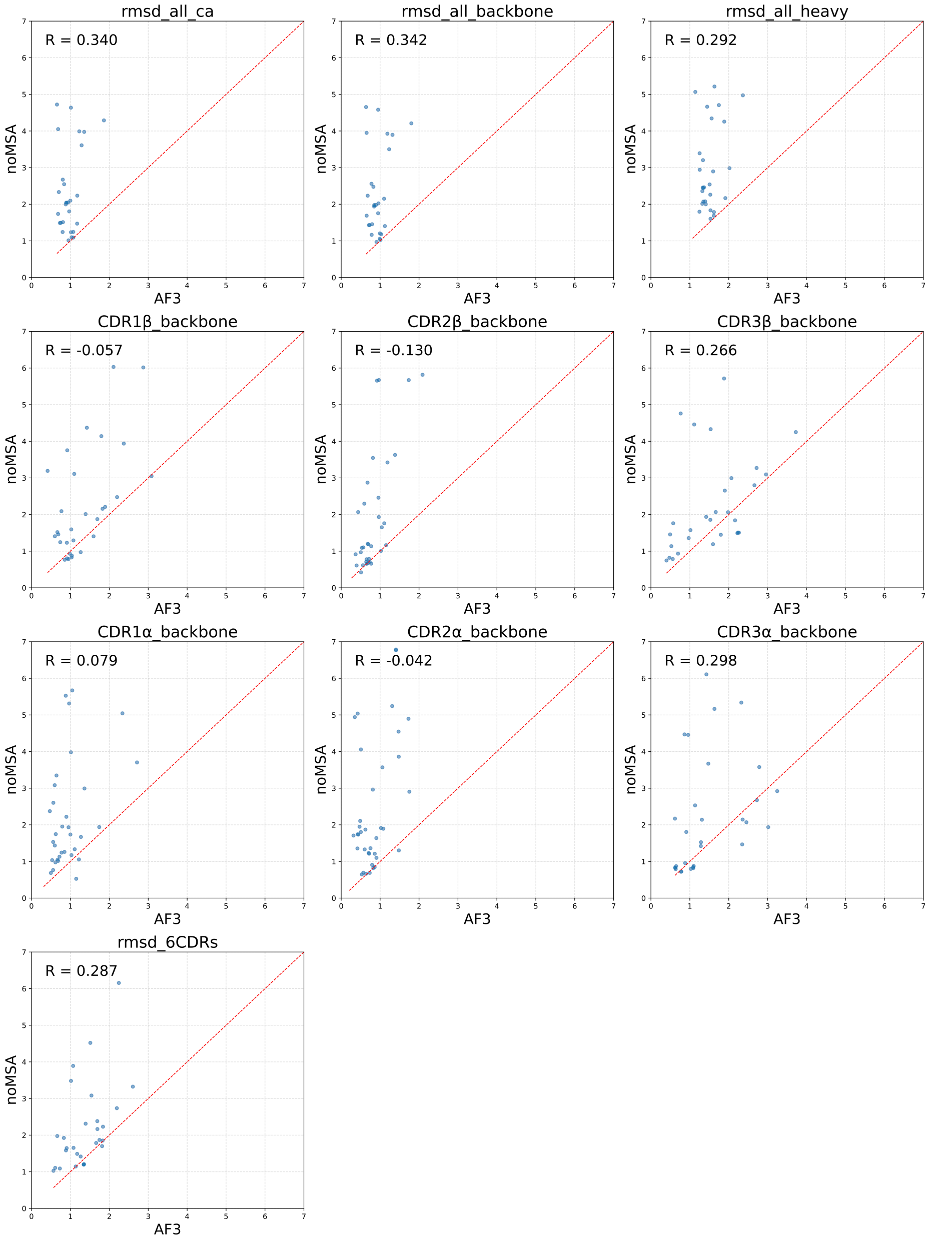


**SI Figure 3.** RMSD correlations for TCR modelling using the native AF3 workflow and one variant using **noMSA**. RMSD were calculated between the experimental structure and the model generated by a given AF3 workflow, on the entire TCR based on heavy atoms (rmsd_all_heavy), on Cα atoms (rmsd_all_Ca), and on backbone atoms (rmsd_backbone), as well as for the backbone of the CDR1β (CDR1β _backbone), CDR2β (CDR1β _backbone), CDR3β (CDR1β _backbone), CDR1a (CDR1a_backbone), CDR2a (CDR2a_backbone), and CDR3a (CDR3a_backbone) loops.

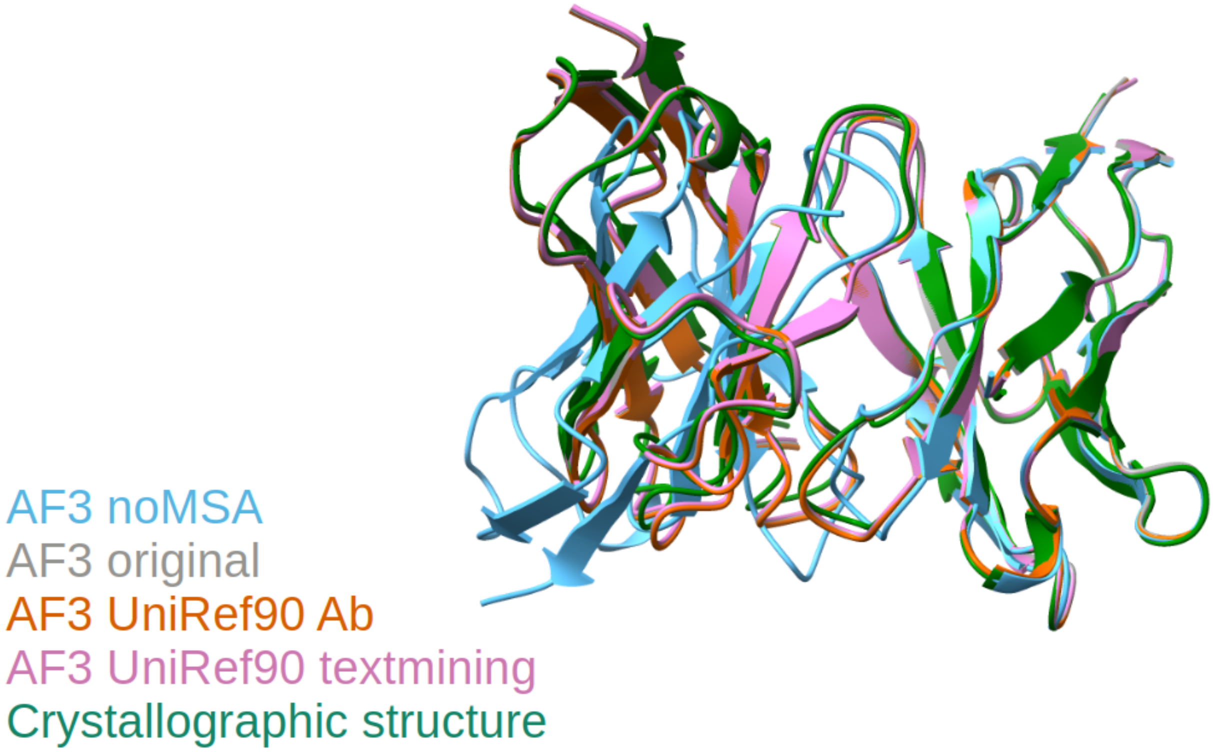


**SI Figure 4.** Super imposition of models and crystallographic structure (pdb : 8zd5) with structural models of the same Ab modelled by AF3 using four AF3 variants: no MSA (in blue), AF3 (in gray), UniRef90-Ab (in orange), and UniRef90-textmining (in violet) superimposed to the crystal structure (in green).

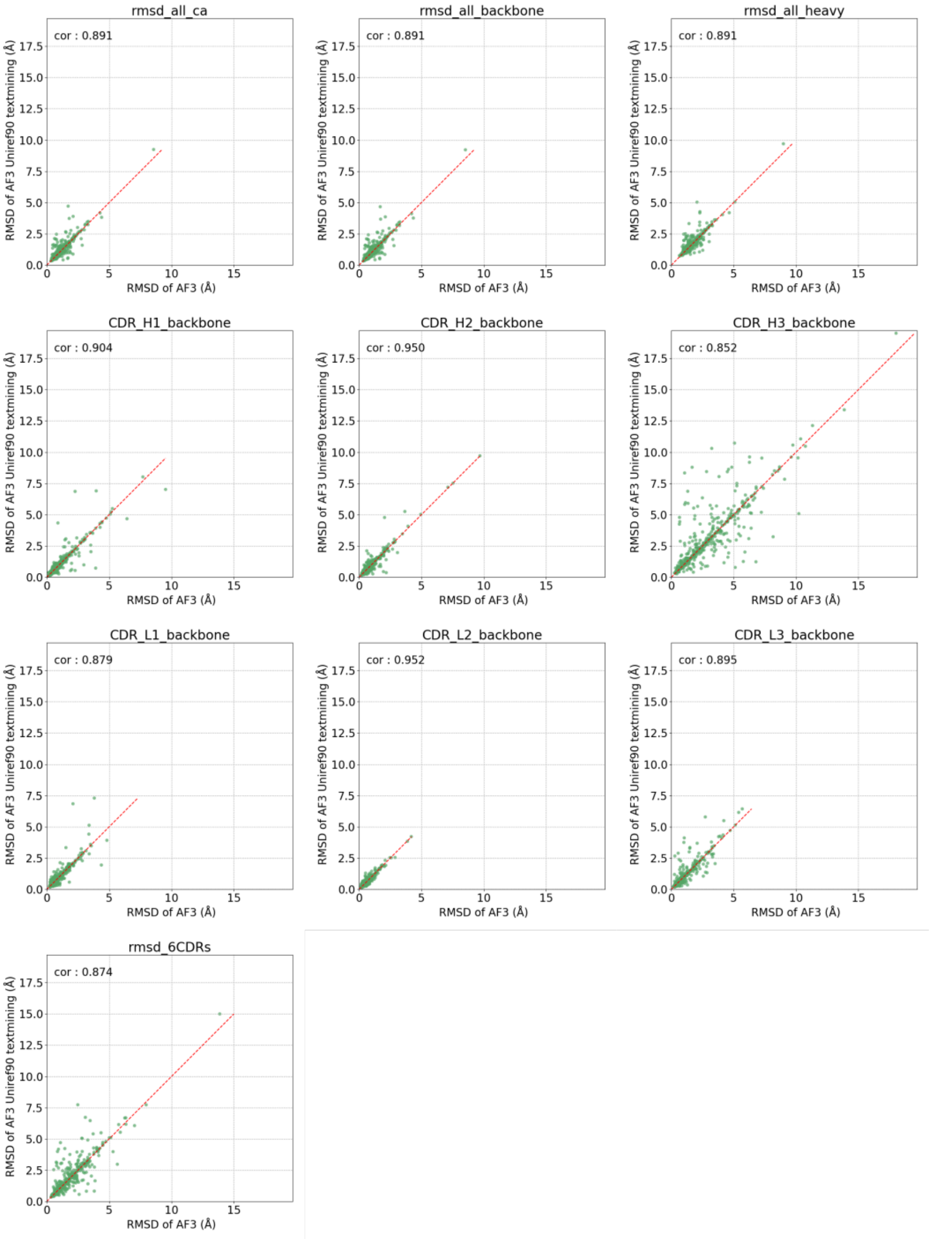


**SI Figure 5.** RMSD correlations for Ab modelling using the native AF3 workflow and one variant using **Uniref-textmining**. RMSD were calculated between the experimental structure and the model generated by a given AF3 workflow, on the entire Ab based on heavy atoms (rmsd_all_heavy), on Cα atoms (rmsd_all_Ca), and on backbone atoms (rmsd_backbone), as well as for the backbone of the CDR H1 (CDR_H1 _backbone), CDR H2 (CDR_H2_backbone), CDR H3 (CDR_3H_backbone), CDR L1 (CDR_L1_backbone), CDR L2 (CDR_L2_backbone), and CDR L3 (CDR_L3_backbone) loops. In red, is a y = x dashed line for reference.

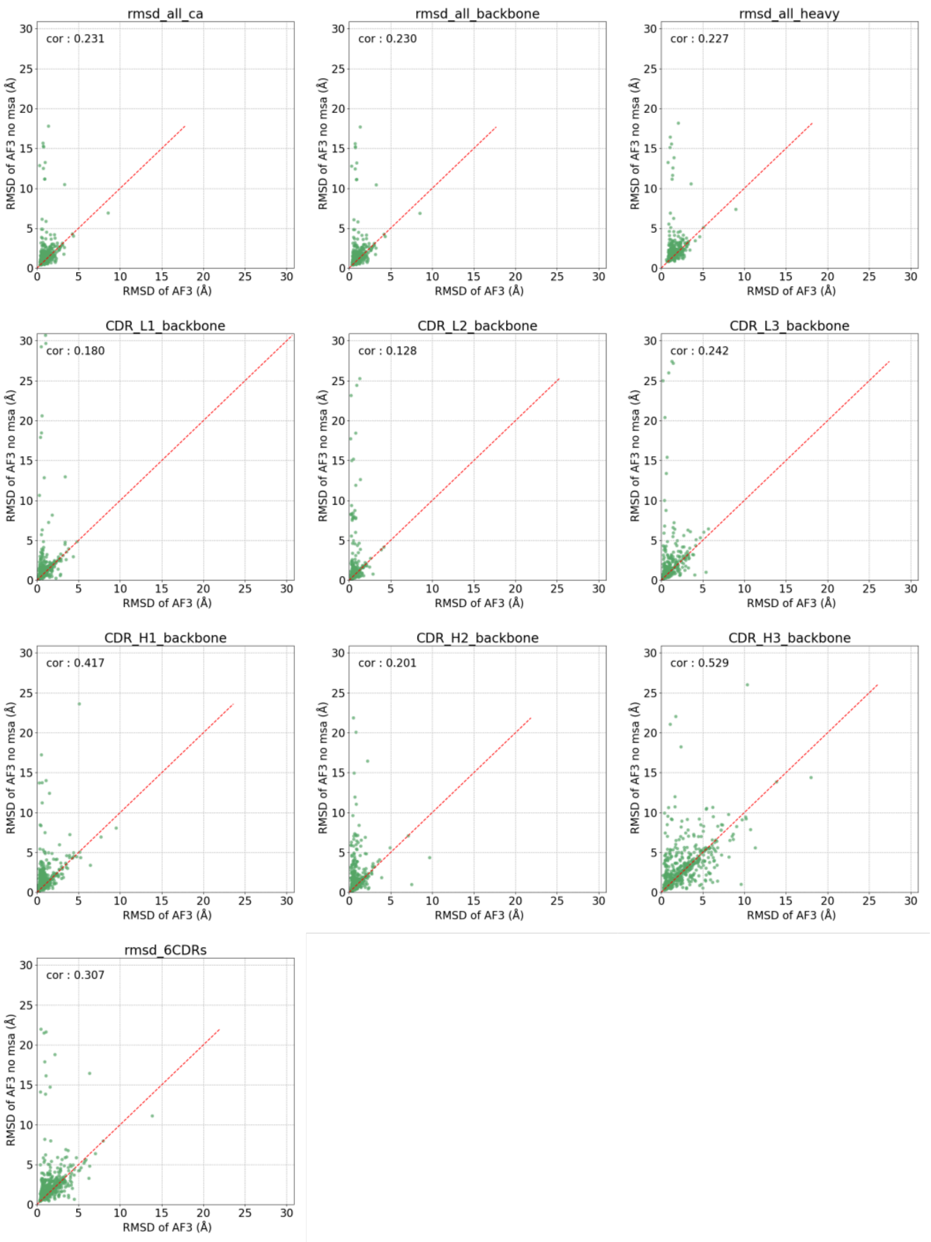


**SI Figure 6.** RMSD correlations for Ab modelling using the native AF3 workflow and one variant using **no MSA**. RMSD were calculated between the experimental structure and the model generated by a given AF3 workflow, on the entire Ab based on heavy atoms (rmsd_all_heavy), on Cα atoms (rmsd_all_Ca), and on backbone atoms (rmsd_backbone), as well as for the backbone of the CDR H1 (CDR_H1 _backbone), CDR H2 (CDR_H2_backbone), CDR H3 (CDR_3H_backbone), CDR L1 (CDR_L1_backbone), CDR L2 (CDR_L2_backbone), and CDR L3 (CDR_L3_backbone) loops. In red, is a y = x dashed line for reference.

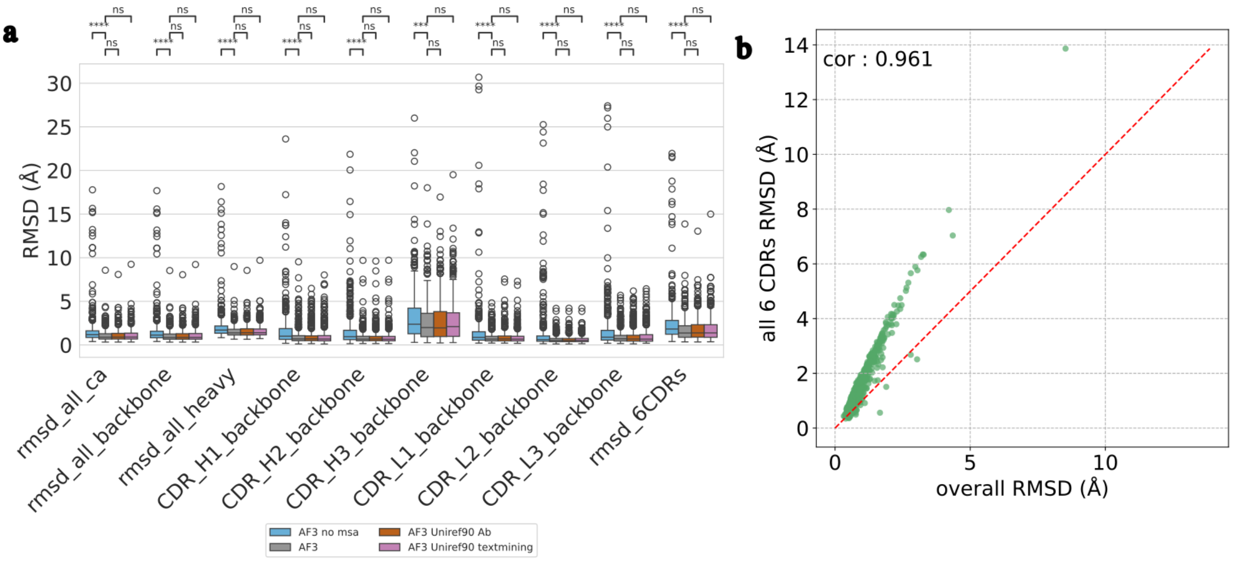


**SI Figure 7**. Complementary information on RMSD analysis of AF3 workflows and on using all 6 CDRs RMSD. **a**) Non-cropped version of RMSD analysis using different AF3 workflow variants for Ab modelling presented in **Figure 6 a)**. **b)** Correlation between the RMSD of all 6 CDRs and the RMSD of the whole structure to ensure that framework residues were routinely very accurately modelled and the deviation in overall RMSD was mostly the reflection of a deviation in CDRs RMSD.

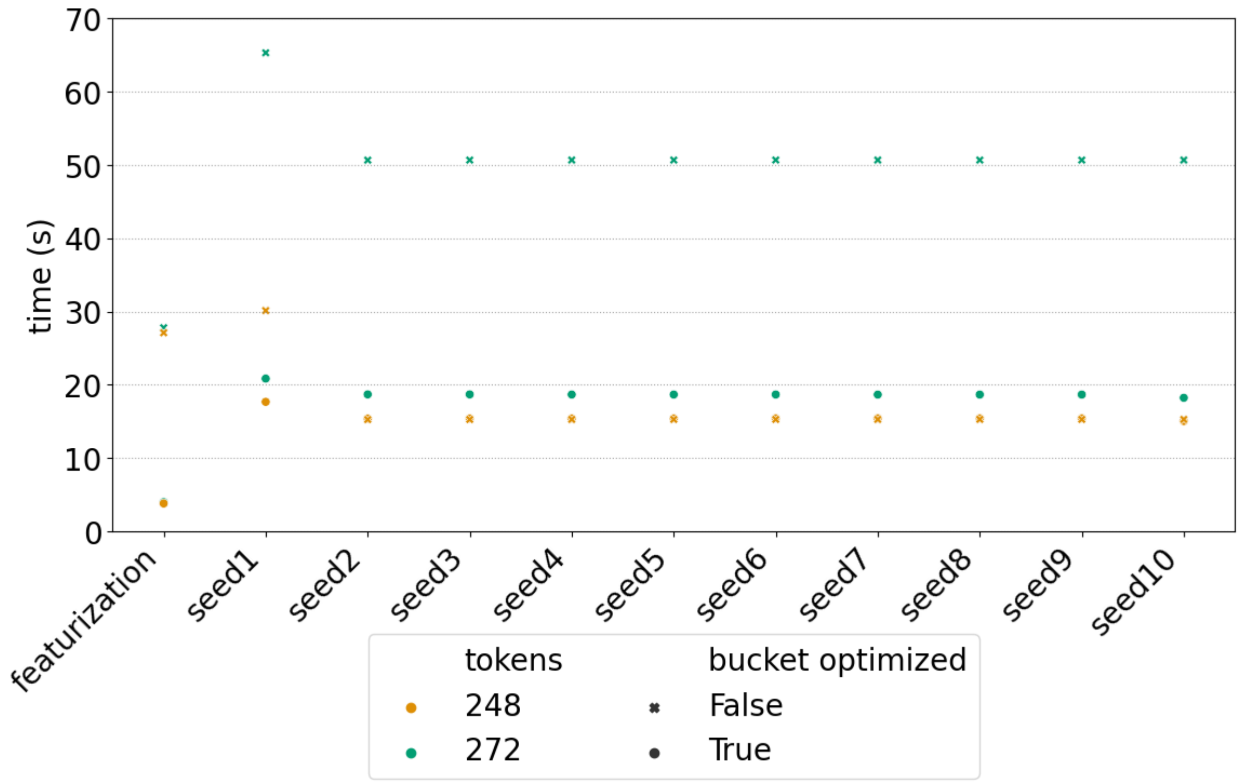


**SI Figure 8**. AF3 inference modelling Ab with and without inference optimization, chosen tokens represent extreme cases of best and worst speed up gains. Inference time for Ab of size 248 tokens with and without bucket optimization overlap between seed 2 and seed 10.
